# Structural and functional characterization of two conserved lumenal TPM-domain proteins involved in the maturation of Photosystem II

**DOI:** 10.1101/2024.10.28.620569

**Authors:** André Vidal-Meireles, Soazig Malesinski, Noémie Lourenco, Yann Horrenberger, Stefania Viola, Caroline L. Monteil, Pascal Arnoux, Corinne Cassier-Chauvat, Jean Alric, Marina Siponen, Xenie Johnson

## Abstract

Many proteins conserved across oxygenic phototrophs play essential roles in photosynthetic function and acclimation, yet many remain unidentified or poorly characterized. In the green alga *Chlamydomonas reinhardtii* we identified two paralogous thylakoid lumenal proteins, encoded by *Cre15*.*g636050* (*LMTP1*) and *Cre03*.*g154600* (*TLP26*), belonging to the conserved TPM-domain (TLP18.3-Psb32-MOLO1) PF04536 family. We generated *C. reinhardtii* knock-out mutants for *LMTP1* and *TLP26* and produced recombinant proteins to assess their biochemical properties. The crystal structures of both proteins reveal the presence of a conserved redox-responsive cysteine pair, novel in TPM-domain proteins, and show that LMTP1 binds manganese. Double mutants lacking both LMTP1 and TLP26 show reduced photosynthetic performance due to effects at the acceptor-side of Photosystem II (PSII); however, these mutants accumulate more chlorophyll and photosynthetic proteins due to increased synthesis rates, a phenotype not observed for the single *lmtp1* and *tlp26* mutants. We propose that these two TPM-domain proteins, LMTP1 and TLP26, are functionally redundant in maturing nascent PSII intermediates during assembly and repair.

## Introduction

Photosynthetic organisms reduce CO_2_ into organic compounds using the sole energy of sunlight. They are the primary producers of organic matter on earth. The photoinduced splitting of water into O_2_ and protons, takes place at the level of the Mn_4_O_5_Ca-cluster of the oxygen-evolving complex (OEC) of photosystem II (PSII), exposed to the thylakoid lumen (1). The production of highly oxidizing reaction intermediates at the donor side of PSII and of reducing compounds at its acceptor side requires many molecular factors for assembling, stabilizing and repairing this protein complex. Structure and function of the photosystems and their arrangement in the thylakoid membrane are constantly adjusting to changes in their environment, especially light, and regulatory mechanisms allow them to maintain performance and to cope with environmental stress. Repair of PSII subunits is a tightly regulated process that, in chloroplasts, involves migration from thylakoid grana to the stromal lamellae, before degradation of the reaction centre D1 subunit and its replacement by a newly synthesized one (2). Different redox-regulated lumenal proteins participate in this process, including proteases – such as FtsH and Deg – and TPM-domain proteins (3, 4). However, many of these components are yet to be identified.

The TPM-domain superfamily of proteins is named after the first three identified proteins containing this domain: **T**LP18.3, **P**sb32, and **M**OLO-1. Formerly known as DUF477, this family of proteins is broadly conserved in prokaryotic and eukaryotic organisms, and is characterized by a conserved secondary structure – a central hydrophobic β-sheet, made of four β-strands, flanked by four amphiphilic α-helices – even though the amino acid sequence between different TPM-domain proteins is rather variable (5, 6). TLP18.3 is involved in D1 turnover during PSII repair, as *Δtlp18*.*3* mutants from the higher plant *Arabidopsis thaliana* (*A. thaliana*) have slower D1 degradation rates (3). The cyanobacterial homologue of TLP18.3, Psb32, also plays a similar role in PSII repair during high light acclimation (4). These proteins are not regarded as essential subunits of PSII, since mutants of those proteins are still viable, but rather as accessory proteins required to confer fitness and aid in the process of acclimation.

Given that all members of the TPM domain family share a similar protein fold (7), it is likely that other TPMs play a role in the regulation of PSII. TLP15 (PTHR35514 / Thylakoid_lumenal_15kDa-like), part of a subfamily of TPM-domain containing proteins, came to our attention because of its conservation across all oxygenic phototrophic organisms and co-evolution with PSII (8). The green algae *Chlamydomonas reinhardtii* (*C. reinhardtii*) encodes two putative paralogues of this family of proteins, one encoded by *Cre15*.*g636050* and previously annotated as CPLD31, and another encoded by *Cre03*.*g154600*, previously annotated as TLP15.2.

Based on the functional results on the single and double mutants of these genes, and on the crystal structures of these proteins, we propose that the two paralogues are distinct proteins, with redundant functions, that evolved from a common cyanobacterial ancestor. We also propose to update the annotation of these proteins to **L**umenal **M**n-binding **T**PM-domain **P**rotein **1** (LMTP1) and **T**hylakoid **L**umenal **P**rotein **26** kDa (TLP26), respectively. These annotations are consistently used throughout the manuscript.

## Results

Genomics is a powerful tool to analyze the origins of oxygenic photosynthesis and predict which genetic factors could be the most relevant to photoprotection and the control of photosynthetic yields. A coarse filtering of all oxygenic phototrophs sequenced to date identified some 85 conserved gene families split into 50 families of genes coding for known protein subunits and regulators of the photosynthetic apparatus, and another 35 gene families with unknown function (9). We applied a fine filter to this latter subgroup (8), by comparing it against the cyanobacterial genomes previously analyzed (12), and searched for conserved genes that have evolved with photosynthesis but were lost in the cyanobacterial accessions that have lost PSII. In this analysis we identified the genes corresponding to *slr0575* and *sll1071* in *Synechocystis* sp. PCC 6803 (*Synechocystis*) as always strictly co-occurring with PSII (Table S1).

### LMTP1 is the paralogue conserved in all oxygenic phototrophs while TLP26 is only present in a cluster of primary endosymbionts

To study the evolution and conservation of Sll1071 across oxygenic phototrophs, we built phylogenetic trees using the maximum likelihood method (Figure S1) of the protein family PTHR35514 (Thylakoid_lumenal_15kDa-like). Given the evolutionary history of photosynthesis (13, 14), it is very likely that this family of proteins in photosynthetic eukaryotes emerged from an endosymbiotic cyanobacterial ancestor, followed by a duplication event in the last common ancestor to all Streptophyta *–* including green algae, mosses, and land plants. We observed that *C. reinhardtii* harbors two paralogous proteins related to Sll1071: **(i)** CrLMTP1 (encoded by *Cre15*.*g636050*) that clusters with proteins of land plants and that is conserved in all oxygenic photosynthetic organisms, and **(ii)** CrTLP26 (encoded by *Cre03*.*g154600*) that clusters with proteins conserved only in some primary endosymbionts. The longer external branches, indicative of higher evolutionary rates, support the neofunctionalization of TLP26 homologues, that we predict to have arisen from a duplication event after the primary endosymbiosis, while the function of LMTP1 homologues likely remained similar to that of the prokaryotic ancestor. We also compared the mature sequences (i.e., without transit peptides as predicted by TargetP 2.0 (15)) of both *C. reinhardtii* paralogues with the sequence of LMTP1 homologues from *A. thaliana* and *Synechocystis*, as well as the sequence of another TPM-domain protein from *C. reinhardtii*, TLP18.3 (Figure S2); these multiple sequence alignments show that CrLMTP1 is the homologous protein that has the strongest sequence similarity with those of both *Synechocystis* and *A. thaliana*, confirming that LMTP1 is the protein conserved throughout all oxygenic phototrophs.

We used DeepTMHMM (16) to predict the secondary structures of both LMTP1 and TLP26 from *C. reinhardtii*; according to this tool, LMTP1 is a soluble protein, while TLP26 contains three C-terminus transmembrane α-helices (Figure S3a). Both CrLMTP1 and CrTLP26 are predicted to be localized in the thylakoid lumen using TargetP 2.0 (15). Previously, using a Venus-tagged construct, LMTP1 was confirmed to co-localize mostly to the chloroplast of *C. reinhardtii* (17), and in *A. thaliana* it was shown to be part of the free fraction of the thylakoid lumen proteome (18). To confirm the targeting of CrTLP26 to the chloroplast and its docking to the thylakoid membrane via the predicted C-terminus transmembrane helices, antibodies against CrLMTP1 and CrTLP26 were raised using recombinant proteins produced in *E. coli*; for LMTP1, the recombinant protein consisted of the full length, mature, protein sequence (i.e. without the predicted N-terminus transit peptide), and for TLP26, the recombinant protein consisted of the soluble domain of TLP26 (i.e without the predicted C-terminus transmembrane α-helices – see Figure S3a); these antibodies were confirmed to specifically recognize their respective paralogue (Figure S3b). We performed immunoblot analyses on either the soluble or membrane-bound proteins from whole cell extracts or thylakoids (using PetA as a marker for chloroplast membrane-bound proteins) and observed that CrTLP26 could only be found in the membrane-associated samples, both from whole-cell extracts and from isolated thylakoids (Figure S3c), thus confirming the secondary structure and localization predictions.

### The crystal structure of recombinant CrLMPT1 and CrTLP26 reveal non-canonical TPM-domain folds and a metal-binding site

Although some progress has been made in the understanding of the cellular role of TPM-containing proteins, the relationship between structure and function is not clear. While several structures have been elucidated, there appears to be no consensus on the role of TPM-domain containing proteins, nor is there a single clearly attributable function to the structural motif that is a TPM-domain. The structure of six TPM domain containing proteins have been determined either by X-ray crystallography or NMR: BA42 from *Bizionia argentinensis* [PDB codes: 2MPB and 4OA3], BA41 from *B. argentinensis* [PDB code: 5ANP], TLP18.3 from *Arabidopsis thaliana* [PDB code: 3PTJ] (6, 19, 20), Rhom172_1776 from *Rhodothermus marinus* [PDB code: 7TBR], CG2496 from *Corynebacterium glutamicum* [PDB code: 2KPT], and PG0361 from *Porphyromonas gingivalis* [PDB code: 2KW7], with the last three being unpublished structures at the time of this submission.

To gain more insights into the molecular function of CrLMTP1 and CrTLP26, we obtained crystals and performed X-ray diffraction experiments, which were complemented with nano-differential scanning fluorimetry (nanoDSF) measurements on the recombinant proteins. In brief, tagged recombinant proteins were purified using affinity chromatography, followed by cleavage of the hexahistidine-tag and size exclusion chromatography (SEC); the purified proteins were used for both nanoDSF experiments and sitting drop vapor diffusion crystallization trials. The later resulted in well diffracting crystals, up to 1.9 Å for both CrLMTP1 and CrTLP26 (Figures 1a and 1b).

**Figure 1.**
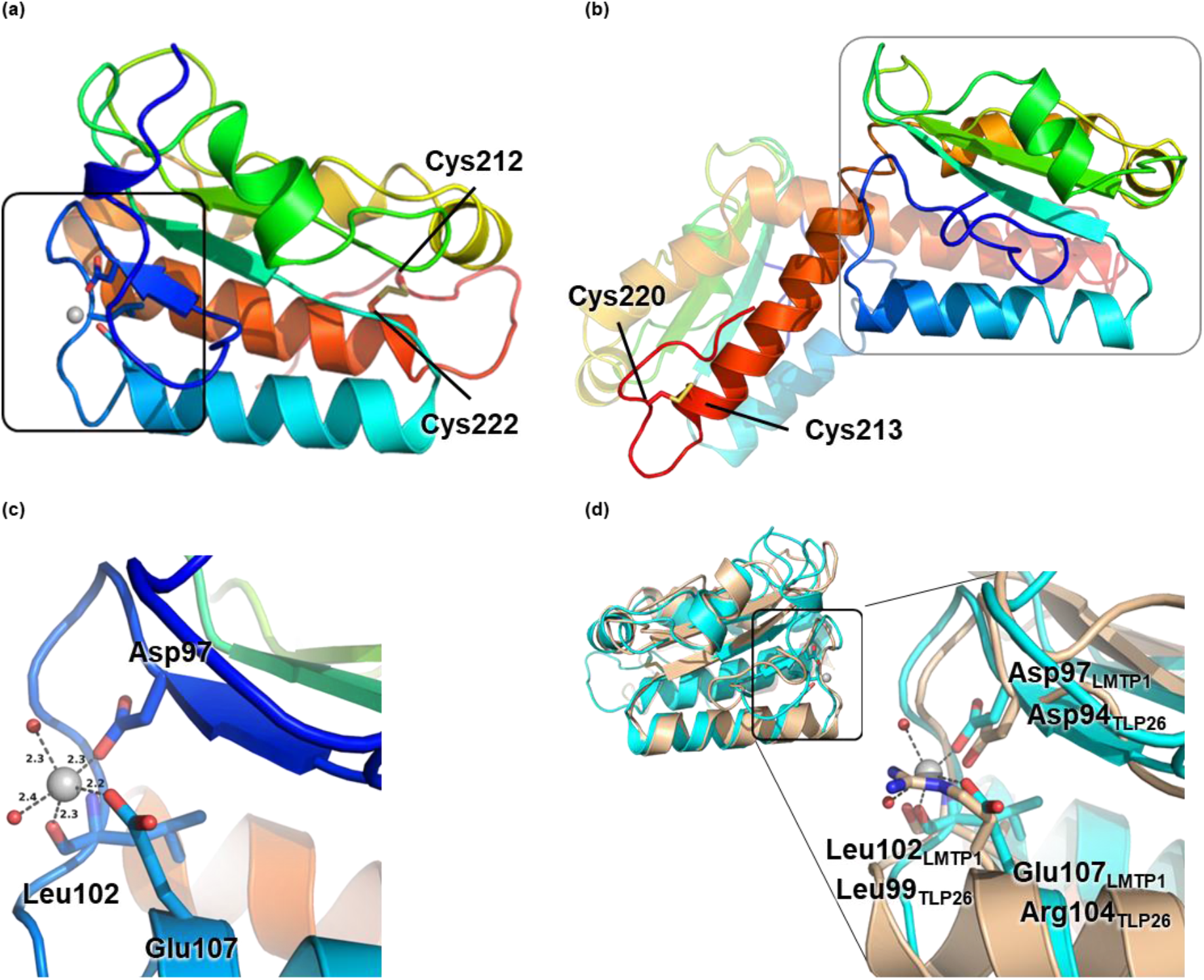
Soluble TPM-domain crystal structures of CrLMTP1 (PDB ID: 9H70) and CrTLP26 (PDB ID: 9TNJ). **(a)** The overall crystal structure of CrLMTP1, solved to 1.9 Å, shows the presence of a near-canonical TPM-domain fold, a metal-binding site at one of the N-terminal loops of the protein (surrounded by a black rectangle), and two cysteine residues forming a disulfide bond at the C-terminus of the TPM-domain. **(b)** The crystal structure of CrTLP26, solved to 1.9 Å, shows the presence of a similar non-canonical TPM-domain fold with the same structural features as the CrLMTP1 protein, including the two cysteine residues forming a disulfide bond at the C-terminus of the TPM-domain, but without the metal binding site; for CrTLP26, however, the fold is obtained through a helix swapped dimer where the C-terminal helix of chain A is swapped with the same helix of chain B. **(c)** The metal bound to CrLMTP1 is hexa-coordinated, and interacting with the residues Asp97, Leu102, and Glu107 (Asp22, Leu26, and Glu32 in the recombinant protein sequence). **(d)** Super-position of CrLMTP1 (cyan) and CrTLP26 (wheat) structures gives a r.m.s.d. value of 1.18 Å^2^ over 130 residues, highlighting the high similarity of the overall TPM-domain fold of both proteins; zooming into the structure of CrTLP26, unlike for CrLMTP1, there is no identifiable metal-binding site due to the differences in protein sequence, namely the substitution of Glu107 from CrLMTP1 into Arg104 in CrTLP26 (Arg24 in the recombinant protein sequence).

In the first case, the CrLMTP1 crystal structure (PDB ID: 9H70) presents with a near canonical TPM fold (Figure 1a) – a central hydrophobic β-sheet, made of four β-strands, flanked by amphiphilic α-helices on each side (6). Three α-helices are present on one side and a single small helix on the other side of the β-sheet which runs from the N-terminus through to the C-terminus with the topology A(↑)B(↑)C(↑)D(↓). α-Helices I, IV, and V are packed against one side of the β-sheet, while α-helix II is located on the opposite side, constituting an αβα sandwich structure. α-Helix III is perpendicular to the plane of the β-sheet, which represents the middle layer of the sandwich. The locations of all these regular secondary structure elements are: β-strands A (residues 95-98), B (125-127), C (169-176), and D (163-167), and α-helices I (102-121), II (138-144), III (179-188), IV (191-196), and V (198-214). A notable difference in the αβα sandwich structure is in the “lack of substance” in the second α-layer of the sandwich which is composed of the very small, 6 amino acid, α-Helix II and loops.

Additionally, the CrLMTP1 structure contains a metal cofactor binding site at one of the N-terminal loops between β-strands A and α-helices I, a region which is generally structurally conserved in other TPM-domain proteins, but which is found empty in other structures. The binding site in CrLMTP1 is occupied by a divalent metal, hexa-coordinated by waters, the side-chains of protein residues Asp97, Glu107, and the carbonyl group of Leu102, which are all conserved in the *Synechocystis* and *A. thaliana* homologues of CrLMTP1, but not in the algal paralogue CrTLP26 nor in TLP18.3 (Figure 1c). The crystal structure also shows a cysteine pair at the C-terminus of the protein that is oxidized, forming an intra-molecular disulfide bridge, which could be important for the function and stability of the protein (Figure 1a).

With no divalent cations present in the crystallization solution, the modelled divalent ion in the LMTP1 structure is Mg^2+^, which best fits the observed density. This cation likely binds in *E. coli* and is not eliminated during our purification protocol. To determine what divalent metal really binds to recombinant CrLMTP1, and to see if the same metal could also bind to TLP26, we used nanoDSF with an array of different divalent metals in a 1:5 (protein:metal) ratio. From all tested divalent metals, only Mn^2+^ (which typically displays hexa-coordination, in agreement with the cofactor binding site of our structure) significantly increased the melting temperature of CrLMTP1 (Figure 2a); for TLP26, no tested divalent metal caused a significant shift in the thermal stability of the protein (Figure 2b).

**Figure 2.**
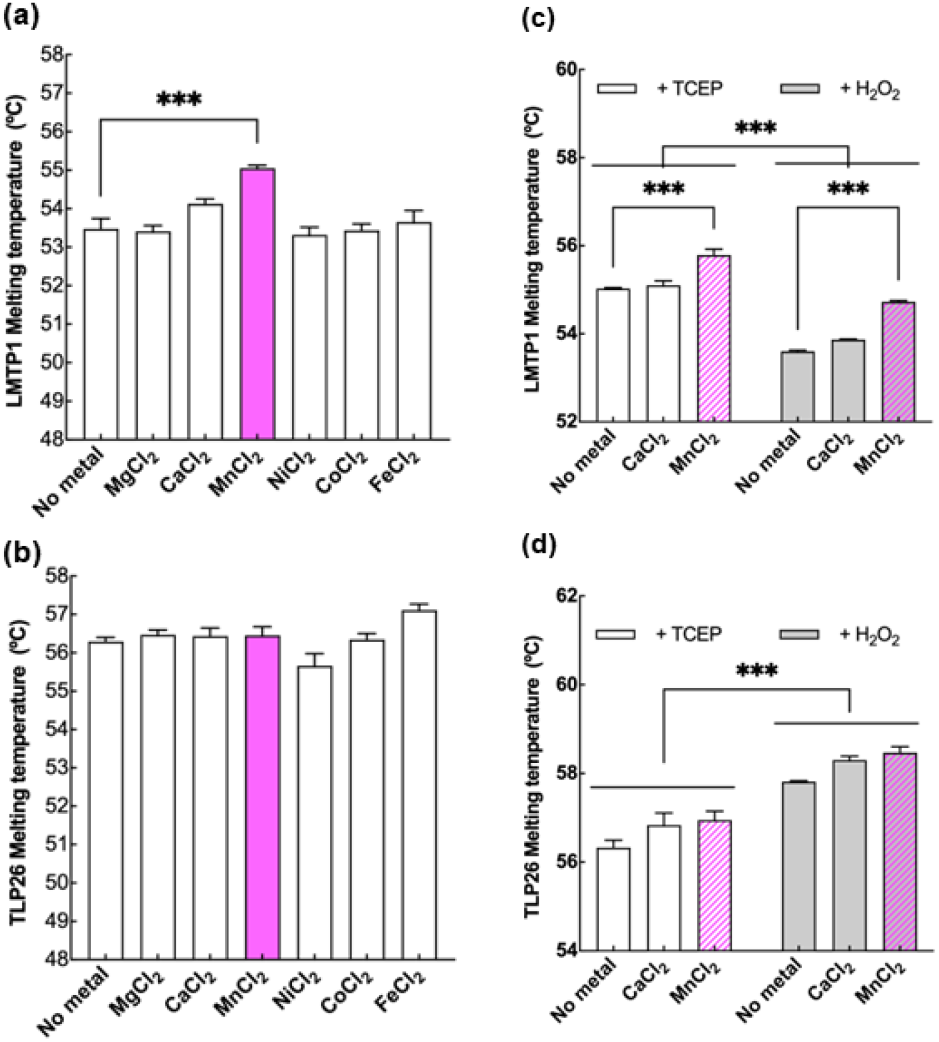
Recombinant CrLMTP1 shows binding to manganese (MnCl_2_), and the melting temperature of both CrLMTP1 and CrTLP26 changes with the redox poise of the medium. NanoDSF thermal denaturation assays for recombinant CrLMTP1 **(a)** and CrTLP26 **(b)**, showing variations in the protein melting temperature (Tm) upon incubation with 5 molar equivalents of different divalent metals. Denaturation assays for CrLMTP1 **(c)** and CrTLP26 **(d)** incubated under reducing (+ TCEP) or oxidizing (+ H_2_O_2_) conditions, in the absence or presence of 5 molar equivalents of the selected divalent metals. Data is represented as mean ± standard error of the mean (n = 3 replicates). *** p<0.001 vs. “No metal” or “+ TCEP”.

The crystal structure of CrTLP26 (PDB ID: 9TNJ) also presents with the same non-canonical TPM fold, as described for CrLMTP1 (Figure 1b). However, this fold is obtained through a helix swapped dimer where the C-terminal helix of chain A is swapped with the same helix of chain B; as we have never observed the dimer in solution, for the time being we conclude the biological assembly to be a monomer. Super-positioning of the CrLMTP1 and CrTLP26 crystal structures shows a structural alignment with an r.m.s.d. value of 1.18 Å^2^ over 130 residues, which indicates a highly conserved fold between these two proteins (Figure 1d). As for our CrLMTP1 structure, CrTLP26 points to the importance of the cysteine pair at the C-terminus of the protein, which is forming an intra-molecular disulfide bridge (Figure 1b).

Since we observed that the recombinant CrLMTP1 and CrTLP26 cysteines enable the formation of intramolecular disulfide bridges, we investigated the effect of the redox status of these residues on the protein conformation (Figures 2c and 2d). We incubated the protein under oxidizing (+ H_2_O_2_) or reducing (+ TCEP) conditions, and in the absence or presence of selected divalent metals. Changing the redox poise of the medium caused shifts in the thermal stability of both recombinant proteins; in the case of CrLMTP1, in both conditions incubation with Mn^2+^ significantly changed the melting temperature, indicating a stabilization of the protein in comparison to the condition with no metals added (Figure 2c).

### *Double* C. reinhardtii *knock-out mutants for LMTP1 and TLP26 show increased PSII maximum quantum efficiencies and synthesis of chlorophyll-binding photosynthetic proteins*

None of the mutants available from the *C. reinhardtii* CLiP library (21) had clear insertions in the coding regions of either of our target genes, so we generated *lmtp1* and *tlp26* mutants via CRISPR/Cas9. The first exons of each gene were targeted using *in vitro* assembled RNP particles. Based on bioinformatics predictions (22, 23), we selected CCTCGGCACGCTGGAAGTTA<u>CGG</u> (located within the first exon of *Cre15*.*g636050*) and CAAATGGCAGCGAGCTACAG<u>CGG</u> (located within the first exon of *Cre03*.*g154600*) as targets. Gene disruption was confirmed via PCR (Figure S4a), and immunoblot analysis, using the antibodies raised against the recombinant LMTP1 and TLP26 proteins (Figure S3b) confirmed the absence of the respective protein in each mutant (Figure S4b).

When the *lmtp1* and *tlp26* single mutants were exposed to illumination with low (20 μmol photons m^-2^ s^-1^) and moderate high light (250 μmol photons m^-2^ s^-1^) under photomixotrophic conditions (i.e. in the presence of acetate as reduced carbon source), both mutants grew at a similar rate as WT cells (Figure S4c). The mutants accumulated equal chlorophyll (*a*+*b*) levels (Figure S4d) and showed maximum PSII quantum yields comparable to that of the WT (Figure S4e).

The immunoblots using the primary antibodies against CrLMTP1 and CrTLP26 showed the accumulation of each protein in the absence of the other in the *lmtp1* and *tlp26* mutants (Figure S4b), suggesting that they are not part of the same complex or regulatory pathway, but not excluding the possibility of a potential overlap in function. To investigate this hypothesis, double *lmtp1 tlp26* mutants were generated by genetic crossing of the two single mutants, and after tetrad separation we obtained several double knock-out mutants that were confirmed to be devoid of both proteins via PCR (Figure S5a) and immunoblot analysis (Figure S5b); three lines were chosen for the following phenotypical tests.

Contrary to the single mutants, we observed a clear phenotype for the *lmtp1 tlp26* lines, that was present in both low and moderate high light, and even at photoinhibitory high light intensities (650 μmol photons m^-2^ s^-1^): chlorophyll (*a*+*b*) levels are significantly increased (Figure 3a). Additionally, at higher light intensities, when the maximum quantum yield of PSII decreased in WT cells due to photoinhibition (Figure 3b), it decreased much less in the double mutants; both phenotypes were not observed in the single mutants (Figures S4d and S4e). Immunoblot analysis showed that the increased chlorophyll contents were due to the accumulation of chlorophyll-binding, photosynthetic antenna proteins, as well as the PSII and PSI reaction centre subunits D1 and PsaA (Figures 3c and S6). To distinguish whether this protein accumulation was due to changes in synthesis or degradation, we exposed the cells to either the chloroplast translation inhibitors lincomycin and chloramphenicol (24) or to the cytosolic translation inhibitor cycloheximide (25) under photoinhibitory high light intensities during 48 h. In the presence of lincomycin and chloramphenicol (“+Linc/Chl”), the *lmtp1 tlp26* lines did not bleach like the WT cells (Figure 3d), even though chloroplast-encoded photosynthetic proteins were degraded at a similar rate as in the WT (Figure S7). This suggested that the nuclear-encoded light harvesting antennas were rather retained in the double-mutants under these conditions. Indeed, in the presence of cycloheximide (“+Chx”), both *lmtp1 tlp26* mutants and WT cells fully bleached after 24 h of illumination (Figure 3d). Altogether, these results indicate that the higher chlorophyll content per cell and accumulation of photosynthetic proteins in the double mutants is due to increased synthesis of photosynthetic-related proteins rather than to a slower degradation.

**Figure 3.**
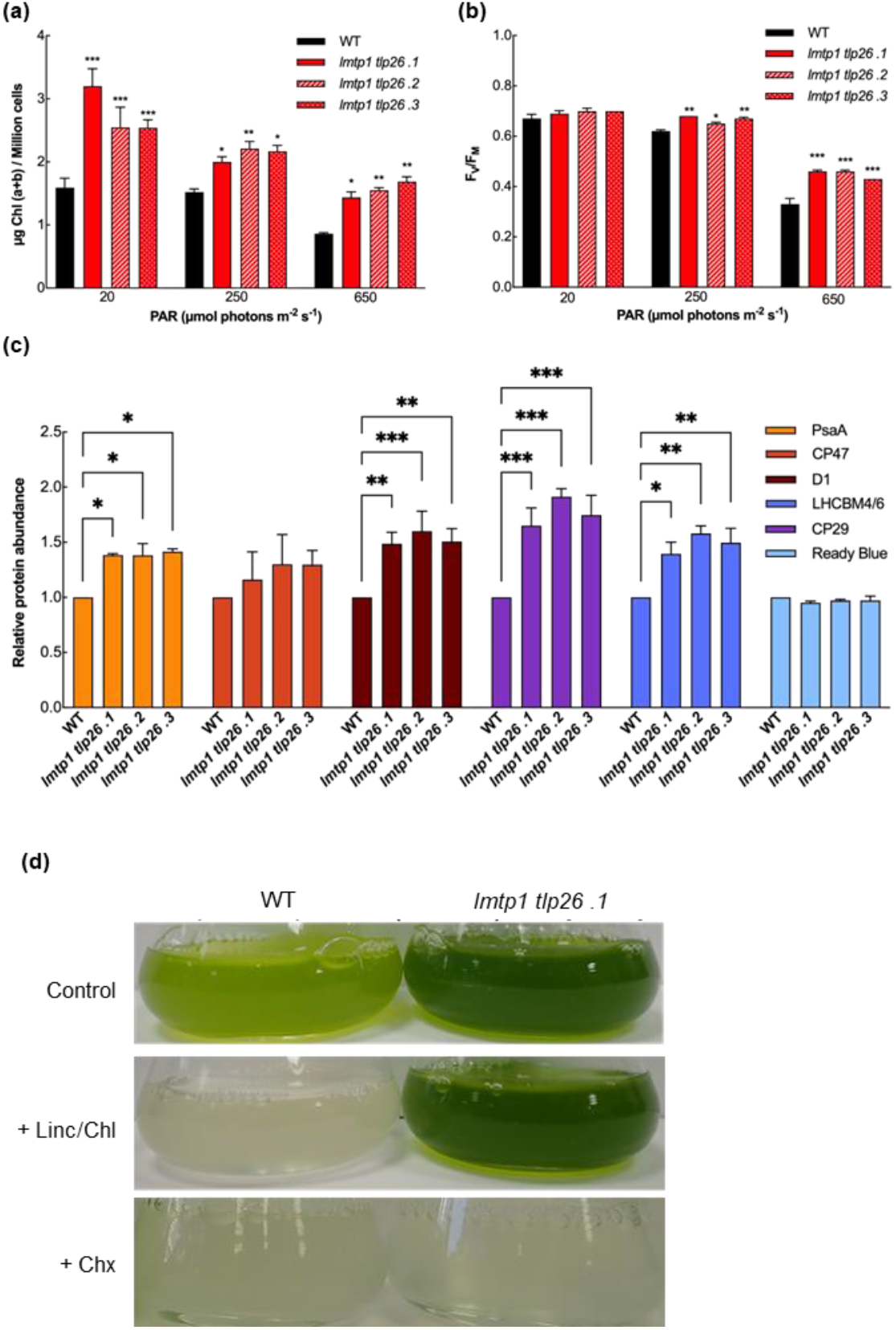
Phenotypical analysis of double *lmtp1 tlp26* mutant lines during growth in photomixotrophic conditions under low (20 µmol photons m^-2^ s^-1^), moderate high (250 µmol photons m^-2^ s^-1^), or photoinhibitory high (650 µmol photons m^-2^ s^-1^) light intensities. **(a)** After 48 h of growth under the different conditions, the double mutants always accumulate significantly more chlorophyll per cell. **(b)** After 48 h of growth at photoinhibitory light intensities, the double mutants and have significantly higher PSII maximum quantum yields (F_V_/F_M_). **(c)** Densitometry analysis of immunoblots (see Figure S6) of membrane proteins from different *C. reinhardtii* strains grown under low light intensities, showing the accumulation of different nuclear and chloroplast encoded photosynthetic subunits; data is normalized for the intensity signal of “100 % (0 h) WT” lane (here presented as “WT”). **(d)** When grown under photoinhibitory light intensities for 48 h, the double mutants continue to accumulate chlorophyll, even in the presence of chloroplast translation inhibitors lincomycin and chloramphenicol (+ Linc/Chl), unless the cytosolic translation of nuclear-encoded genes is blocked by adding cycloheximide (+ Chx). Data is represented as mean ± standard error of the mean (n = 3-5 replicates). * p<0.05; ** p<0.01; *** p<0.001 vs. “WT”.

### The combined absence of LMTP1 and TLP26 causes a decrease in photosynthetic performance due to limitations at the level of the PSII acceptor-side

The functional consequences of the absence of CrLMTP1 and CrTLP26 were further studied on low light grown cells, where the higher chlorophyll phenotype is constitutive but not accentuated by high light treatments. We used dark-induced relaxation kinetics (DIRK) of ElectroChromic Shift (ECS) signals to quantify the total electron transport rates via both photosystems (ETR_I+II_) in the mutant and WT cells in response to increasing light intensities (26): *lmtp1 tlp26* mutants have lower ETR_I+II_ values (Figure 4a), indicating that electron transport across the intersystem chain is compromised. In agreement with these findings, we also observed that CO_2_ fixation rates were lower in the mutants compared to WT cells (Figure 4b).

**Figure 4.**
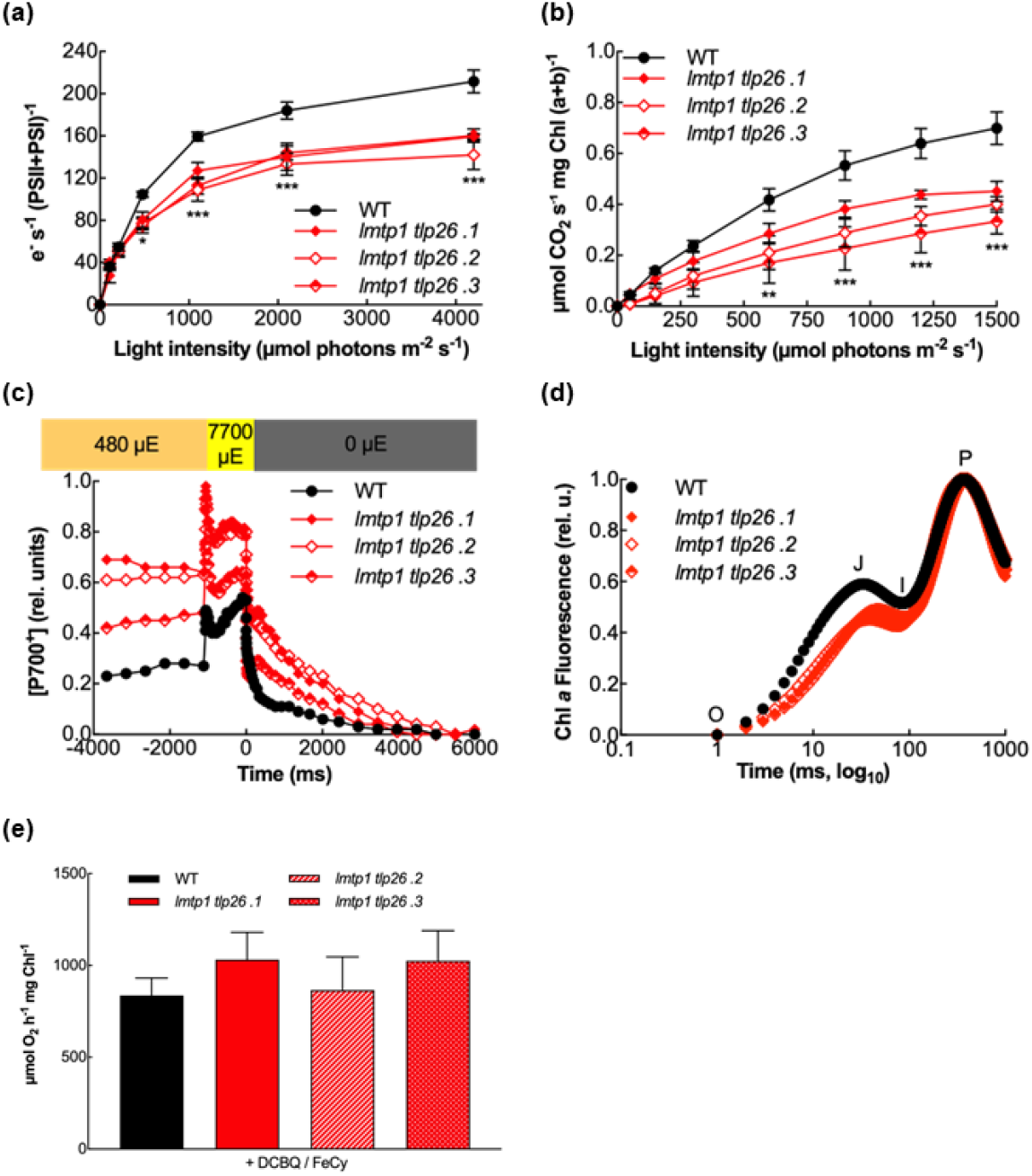
Double *lmtp1 tlp26* mutant lines are constitutively acceptor-side limited at the level of PSII. **(a)** Total electron transport rates (ETR_I+II_) based on DIRK measurements and normalized to the total amount of PSI+PSII reaction centers, as a function of incident light. **(b)** CO_2_ assimilation rates as a function of incident light. **(c)** Representative P_700_ redox kinetics of cells acclimated for 3 min at 480 μmol photons m^-2^ s^-1^ (μE), followed by a 1 s saturating pulse and a dark-relaxation phase; traces are normalized on the maximal amounts of oxidizable P_700_ measured in the presence of DCMU and DBMIB. **(d)** Representative OJIP fluorescence traces of cells grown under low light intensities. **(e)** Maximum O_2_ evolution rates measured in the presence of the artificial electron acceptors DCBQ and FeCy. Data in panels (a), (b) and (e) is represented as mean ± standard error of the mean (n = 3 replicates). * p<0.05; ** p<0.01; *** p<0.001 vs. “WT” at each light intensity.

We monitored P_700_ redox kinetics to estimate the rate of donor- (Y_ND_) or acceptor-side (Y_NA_) limitations at the level of PSI and observed higher donor-side limitations for the *lmtp1 tlp26* lines (Figures 4c and S8a), suggesting a restriction in electron transport upstream of PSI. Fast chlorophyll *a* fluorescence transients (OJIP) showed that F_J_ values were lower in the *lmtp1 tlp26* lines, indicating that the plastoquinone-pool (PQ-pool) was more oxidized in the mutants than in WT cells (Figures 4d and S8b). Altogether, these observations lead to the conclusion that the decreased ETR_I+II_ in the mutants was due to a restriction of electron transport at the level of PSII, even if the synthesis of PSII-related proteins was up-regulated.

To distinguish between donor- and acceptor-side limitations in PSII, we measured maximal O_2_-evolution rates, in the presence of the artificial PSII electron acceptor 2,6-dichloro-1,4-benzoquinone (DCBQ) and potassium ferricyanide (FeCy). Upon a 5 min incubation period at saturating light intensities, the rate of O_2_ evolution was similar in *lmtp1 tlp26* mutants and WT cells (Figure 4e), which indicates that oxygen-evolution (donor side) is not affected in the *lmtp1 tlp26* lines, and thus that the decreased ETR_I+II_ rates must arise from an impairment at the acceptor-side of PSII.

## Discussion

The TPM-domain superfamily of proteins (InterPro entry IPR007621) is widely conserved across bacteria and is limited to invertebrates and phototrophs amongst eukaryotes. The given name “TPM” is based on the first three proteins studied from this family: TLP18.3 from *A. thaliana*, Psb32 from *Synechocystis*, and MOLO-1 from the worm *Caenorhabditis elegans* (7). Despite the low level of conservation in the primary sequences of TPM domain-containing proteins, they can be identified by their conserved and unique secondary structure, composed of a central hydrophobic β-sheet, made of four β-strands, flanked by four amphiphilic α-helices (6). While the precise function and role of most TPM-domain proteins is unknown, as well as the structure-function relationship associated to their physiological role (7), members of this protein family have been proposed to play a role in PSII repair and acclimation to high light in photosynthetic organisms (3, 4).

The uncharacterized family of TPM-domain proteins belonging to the family PTHR35514 (Thylakoid_lumenal_15kDa-like), were included in the GreenCut library published with the *C. reinhardtii* genome, highlighting their conservation in photosynthesis (8, 10). Peptides of PTHR35514 family members have previously been detected in proteomics studies for *C. reinhardtii* (27), *A. thaliana* (18), and cyanobacteria (28); in *A. thaliana*, the AT5G52970 protein was proposed to be a constituent of the soluble fraction of the thylakoid lumen proteome (18), and in cyanobacteria Sll1071 peptides are found in the interactome of the PsbU lumenal subunit of PSII (28). The green algae *C. reinhardtii* encodes two paralogues LMTP1 (encoded by *Cre15*.*g636050*) and TLP26 (encoded by *Cre03*.*g154600*); based on TargetP 2.0 predictions (15), both proteins are addressed to the thylakoid lumen, and according to their predicted secondary structure LMTP1 is a soluble protein, while TLP26 is anchored to the membrane by three C-terminus transmembrane alpha-helices. The sub-cellular, chloroplastic, localization of LMTP1 had already been previously shown for *C. reinhardtii* (17), and in *A. thaliana* it was further confirmed to be a soluble protein that is present in the thylakoid lumen (18), but no information regarding CrTLP26 was available in the literature. By fractionating soluble and membrane proteins from whole cell extracts, we were able to confirm the secondary structure predictions that proposed CrTLP26 to be a membrane-docked protein (Figure S3c). Additionally, the immunoblot analysis on isolated thylakoids also confirmed the targeting of this protein to the chloroplast of *C. reinhardtii* (Figure S3c).

To understand the evolutionary history of LMTP1 and TLP26, and to correctly annotate them, we have compared them to other members of the PTHR35514 family from various prokaryotic and eukaryotic oxygenic phototrophs and built phylogenetic trees (Figure S1). We identified two clear groups of proteins, one containing TLP26 from *C. reinhardtii* and homologues from primary Chlorophyte endosymbionts, and another containing LMTP1 from *C. reinhardtii*, the cyanobacterial homologue Sll1071, and the *A. thaliana* homologue AT5G52970. This division is coherent with the sequence alignment of the different homologues, because TLP26 is the most divergent (Figure S2). Data also suggest that both proteins evolved from a common cyanobacterial ancestor, which was duplicated in the early endosymbiotic species, followed by the specialization of each paralogue. Why plants and algal secondary plastid endosymbionts do not have TLP26 homologues is intriguing and may hold the key to understanding the functional specialization of these paralogues.

The high-resolution crystal structures we obtained for CrLMTP1 and CrTLP26 (Figures 1a and 1b) also point to functional diversification, since while both have a near canonical TPM-domain fold, only CrLMTP1 shows a novel metal-binding site (Figure 1c); this site is coordinated by water molecules and by 3 residues (Asp97, Glu107, and Leu102) that are conserved in LMTP1 homologues from different species. Glu107 is replaced by an arginine in CrTLP26 and TLP18.3 (Figure S2), suggesting the absence of this metal binding site in these proteins, an observation that was confirmed in our crystal structure for CrTLP26 (Figure 1d). Of the six crystal structures available in the PDB for TPM-domain containing proteins, three appear as metal binding proteins: Rhom172_1776, AtTLP18.3, and BA42. Notably, the metal-binding site of CrLMTP1 does not correspond to the proposed Ca^2+^-binding site in AtTLP18.3 (19) nor to the Ca^2+^/ Mg^2+^ sites in BA42 (7, 20), but Rhom172_1776 contains a proposed Mg^2+^ binding site with the same metal coordination sphere as the site we identified in CrLMTP1. Unfortunately, Rhom172_1776 has been deposited without being published so far, making it difficult to infer a function uniquely based on this structure. It is worth mentioning that in both cases the crystallization solution did not contain any divalent cations, suggesting the site has a significant affinity for the metal (as it is still present after all protein purification steps).

Based on nanoDSF assays (Figures 2a and 2b), the most likely metal occupying the cofactor site at CrLMTP1 is Mn^2+^; this is also a novel finding within the TPM-domain family of proteins, as Mg^2+^ and Ca^2+^ have been the only divalent metals reported as cofactors of BA42 and TLP18.3 (7, 19). Interestingly, Mn^2+^ was suggested to play a role in the activity of another TPM-domain containing protein, BA41, but no direct proof of metal binding was shown in that work (6). Mn^2+^ typically binds to hard atoms such as oxygen (O) and nitrogen (N), with a hexa-coordinated geometry as observed in our crystal structure. The amino acids involved in this binding mainly include aspartic acid (Asp) and glutamic acid (Glu), through the oxygen atoms of their side chains (29). Bioinformatic analysis using the *MeBiPred* server (30) confirms that the cation coordination distances observed in our CrLMTP1 structure correspond to those of Mn^2+^ (Figure S9); these observations further support that Mn^2+^ is the cofactor of LMTP1 in the thylakoid lumen, at its physiological context; while the specificity of the observed interaction and the dissociation constant between metal and protein remain to be determined.

The crystal structure of CrLMTP1 and CrTLP26 also revealed that the cysteines at the C-terminus of both proteins form disulfide bonds (Figures 1a and 1b). LMTP1 has been proposed as a target of thioredoxins (31), that can regulate the function of target proteins by changing the redox status of their cysteines. For lumenal proteins, this likely involves long distance transduction of redox signals from the stroma (32). In *A. thaliana*, nearly half of all lumenal proteins have been identified as targets of thioredoxin in *in vitro* studies; this contrasts with the chloroplast stroma, where only about 2% of the proteins are regulated in this manner (33, 34). Examples of redox regulation in the lumen include several proteins that are essential for light acclimation, such as STN7, FKBP13, and VDE (35–38). Besides thioredoxin regulation, proteins also respond to lumen acidification upon high light exposure. The changes in the lumenal pH trigger shifts in the protein redox potentials, and also impact the protonation of different protein residues, which promote conformational changes that shift the accessibility of these residues in the protein, including cysteine pairs (32). Our results show that the redox poise of the medium affects the melting temperature of both CrLMTP1 and CrTLP26 (Figures 2c and 2d), suggesting that changes in the redox status of their cysteines might provide a mechanism to regulate their function.

We also noted that the distance between cysteines depends on the presence or absence of transmembrane helices: for CrLMTP1 and AtLMTP1 (soluble proteins), the distance between these amino acids is longer than for CrTLP26 and Sll1071 (membrane-docked proteins). As there are no cysteines in TLP18.3, these are clearly not a requirement for the formation of the TPM-domain and may rather play a specific role on LMTP1 and TLP26 regulation. Additionally, the transmembrane domains of both CrTLP26 and Sll1071 share some degree of similarity. The C-terminus of CrLMTP1, seems to have retained some homology with the start of the transmembrane region of Sll1071 (Figure S2), supporting the former evolved from the latter after gene duplication and cleavage.

Despite these structural differences, however, LMTP1 and TLP26 seem to have at least partially redundant physiological functions in *C. reinhardtii*. Indeed, while the single *lmtp1* and *tlp26* mutants showed no observable phenotype (Figure S4), the double *lmtp1 tlp26* mutant lines showed a constitutive over-accumulation of chlorophyll (Figure 3a) and of antenna (CP29, LHCBM4/6) and core subunits of PSI (PsaA) and PSII (D1) (Figure 3c), likely due to increased synthesis of these proteins rather than improved stability (Figure S7). Despite larger amounts of PSII- and PSI-related subunits, *lmtp1 tlp26* mutants showed lower CO_2_ fixation rates and impaired electron transport at the acceptor-side of PSII (Figure 4). This suggests that the increased synthesis of photosynthetic proteins may represent a compensatory mechanism or reflect a change in the chloroplast redox homeostasis. The more oxidized redox state of the PQ pool observed in the double *lmtp1 tlp26* mutant (Figure 4d) is consistent with a release of the retrograde repression of chlorophyll binding proteins (39). This, in turn, accounts for the strong “stay-green” phenotype observed during incubation under photoinhibitory light intensities (Figure 3d).

That lumenal proteins affect the PSII acceptor side is hardly unexpected. Mutations on the donor side of PSII (in the lumenal loops of the D1 or D2 core proteins) have likewise been shown to modify the midpoint potentials of the quinones Q_A_ and Q_B_, at its acceptor-side (40, 41). Similarly, the presence or absence of an intact OEC at the lumenal side of PSII also affects the midpoint potential of Q_A_ (42). Our results show that PSII possesses an intact OEC in the *lmtp1 tlp26* double mutants (Figure 4e), indicating that CrLMTP1 and CrTLP26 are not required for its biogenesis. The two proteins could, however, affect the acceptor side of PSII by interacting with its lumenal side during the maturation of assembly intermediates, either via the manganese bound to CrLMTP1 or the conserved cysteine pair present in both proteins.

CrLMTP1 accumulates in the absence of CrTLP26, and reciprocally CrTLP26 accumulates in the absence of CrLMTP1 (Figure S4b). This is in favor of them not being in direct interaction but involved in functionally overlapping pathways. Other proteins of unknown function, like the PSBP-domain proteins (PPDs) (43), could partially compensate for the loss of either CrLMTP1 or CrTLP26 in the single mutants. This hypothesis seems very likely, at least for LMTP1, where yeast-two-hybrid assays have demonstrated that it can interact with CYP38 (44), a thylakoid lumen immunophilin involved in PSII maintenance and proper D1 turnover (45, 46); CYP38 could thus bring LMTP1 close to D1, promoting the interaction between the two proteins even when TLP26 is absent.

Although our data does not point to a direct involvement of CrLMTP1 and CrTLP26 in PSII biogenesis and repair, and the exact molecular function of both proteins remains to be fully elucidated, our results provide evidence for the direct involvement of both paralogues in the maintenance of PSII function, in line with their strict co-occurrence across oxygenic phototrophs. Our results also suggest the participation of LMTP1 and TLP26 in a broader redox network, possibly via their cysteine residues and manganese-binding properties, that contribute to the maintenance of chloroplast redox homeostasis from the thylakoid lumen space.

## Materials and Methods

### Algal cell culture

The *Chlamydomonas reinhardtii* wild-type (WT) strain, T222 (mt+), used in this study is a progeny of 137c backcrosses (47). Maintained cultures were always cultivated under photomixotrophic conditions, at 20 μmol photons m^-2^ s^-1^ in Tris-Acetate-Phosphate (TAP) + 2 % agar media plates. For experimental setups, liquid cell cultures were grown in TAP medium at 25 °C under ambient air at continuous 20 μmol photons m^-2^ s^-1^ illumination in incubation shakers (photomixotrophic conditions) for 3 days, before scaling-up and moving them into the experimental illumination conditions. Standard recipes for media preparation were used as in (48). Experiments were performed using batch cultures in 100- or 500-ml Erlenmeyer flasks. Cell densities were calculated using a Beckman Coulter Z2 instrument and chlorophyll content was determined via methanol extraction (49). For blocking chloroplast protein synthesis, 0.5 mg ml^-1^ of lincomycin and 0.125 mg ml^-1^ chloramphenicol were added to the cells and allowed to incubate for 1 h at 20 μmol photons m^-2^ s^-1^ before placing the cells under 650 μmol photons m^-2^ s^-1^ illumination; for blocking nuclear protein synthesis, 10 µg ml^-1^ of cycloheximide were added to the cells and allowed to incubate for 1 h at 20 μmol photons m^-2^ s^-1^ before placing the cells under 650 μmol photons m^-2^ s^-1^ illumination.

### *Generation of* C. reinhardtii *CRISPR/Cas9 mutants*

The transformation was performed based on (50, 51), with some modifications. Briefly, specific gRNAs and Alt-R^TM^ S.p. Cas9 Nuclease V3 were ordered from IDT-DNA and used to assemble *in vitro* RNP particles. *C. reinhardtii* wild-type cells were harvested at early exponential stage (2×10^6^ cells ml^-1^) via centrifugation (1600 g, 5 min, RT) and re-suspended in “MAX Efficiency™ Transformation Reagent for Algae” (Thermo Fisher Scientific), supplemented with 20 % sucrose, to a final concentration of 2×10^8^ cells ml^-1^. A heat-shock (40 ºC, 30 min, 450 rpm shaking) was then applied to the cells, followed by a 20-min recovery at RT, before 120 µl of cells were added to a pre-cooled 2-mm gap electroporation cuvette together with 15 µl of RNP particles and 1 µg of paromomycin-resistance plasmid for electroporation (600 V, 50 µF, infinite external resistance). Cells were allowed to recover overnight in TAP medium supplemented with 20 % sucrose and 1 mg l^-1^ Vitamin B12 at 33 ºC in the dark before being plated in TAP + 2 % agar + 10 mg l^-1^ paromomycin and incubated under 20 μmol photons m^-2^ s^-1^ illumination until colonies could be picked. Single mutants *lmtp1 mt-* and *tlp26 mt+* were then crossed as described in (52) and progenies were selected via PCR and immunoblot analysis.

### DNA isolation and PCR

Total genomic DNA and PCR were done using the “Phire Plant Direct PCR Master Mix” kit (Thermo Fisher Scientific) according to the manufacturer’s instructions. Briefly, a small amount of cells was solubilized with 20 µl of “Dilution buffer” (included in the PCR kit) *via* agitation, before centrifugation (4000 g, 10 min, RT) to obtain the DNA in the soluble fraction. The PCR master mix was prepared by mixing 5 µl of 2X reagent with 3 µl of 3 M betaine, 1 µl of each primer (10 µM), and 1 µl of DNA template (or ddH_2_O as a negative control), and amplification was done during 35 cycles. PCR products were loaded on a 2 % TAE-agarose gel, and the DNA was stained for visualization with ClearSight (Euromedex); O’GeneRuler Express DNA ladder (Thermo Scientific) was used to estimate amplicon sizes. The primers used to amplify the *lmtp1* gene are P1, 5’-TTGTCAACATGCAGAGCCTC-3’ and P2, 5’-TAGTTGCAGCCAGTAGTCGT-3’ (corresponding to an expected WT-sized amplicon of 249 bp) and for the *tlp26* gene are P3, 5’-ATGCAAGTTTACGGGAATGCA-3’ and P4, 5’-GATGCCAGTACACCACGTCTT-3’ (corresponding to an expected WT-sized amplicon of 748 bp).

### Protein isolation

Protein extraction from whole cell extracts was based on (53) with minor modifications. Briefly, 15-ml of culture were collected at each time-point, centrifuged (20800 g, 1 min, RT), and the pellets were frozen in liquid N_2_ before being re-suspended in 350 µl of lysis buffer (50 mM Tris pH 6.8, 2 % SDS, 1 mM DTT, 1 mM PMSF, 1X cOmplete protease inhibitor without EDTA – Sigma Aldrich) and incubated at 37 ºC for 30 min with strong agitation. Protein concentration was calculated against a bovine serum albumin (BSA) standard curve, using the Pierce™ BCA Protein Assay Kit (Thermo Fisher Scientific) according to the manufacturer’s instructions.

Separation of soluble and membrane-bound protein fractions was based on (54) with modifications. Briefly, 5-ml of culture were collected via centrifugation (1600 g, 5 min, RT), re-suspended in 350 µl of lysis buffer (10 mM Tris pH 8.1, 1 mM EDTA, 1X cOmplete protease inhibitor without EDTA – Sigma Aldrich) and flash-frozen in liquid N_2_, before being thawed at 37 °C and once again flash-frozen in liquid N_2_. After 3 cycles of freeze-thawing, samples were centrifuged (20800 g, 30 min, 4 °C) and the supernatant was stored as the protein soluble fraction; the remaining pellet was then processed for protein extraction as for whole cell extracts and stored as membrane-bound fraction.

### Thylakoid isolation

Thylakoid extraction was done according to the standard method (55) and the thylakoids were resuspended in a final 5 mM HEPES/KOH pH 7.5 at 1 mg ml^-1^ chlorophyll for subsequent immunoblot analysis.

### Immunoblot analysis

Western blot analysis was performed as in (8) with minor modifications. Briefly, proteins were separated under denaturing conditions on 13 % SDS-PAGE gels. Samples were loaded based on total protein content (100 % equals 2 μg protein); loading was confirmed via staining of the SDS-gel with ReadyBlue™ Protein Gel Staining (Sigma-Aldrich) according to the manufacturer’s instructions. Proteins were transferred onto nitrocellulose membranes (BioTrace NT, Pall Corporation) using liquid transfer. Primary LMTP1 and TLP26 antibodies were generated in rabbit against the recombinant protein (ProteoGenix) and used at a 1:5000 ratio; primary CP47, D1 (PsbA), and PetA antibodies were obtained from Agrisera (AS04 038, AS05 084, and AS06 119) and used at a 1:2500, 1:20000, and 1:40000 ratio, respectively; primary PsaA, LHCBM4/6, and CP29 antibodies were a kind gift from S. Bujaldon and used at a 1:10000, 1:10000, and 1:5000 ratio, respectively. Secondary antibodies used were always HRP-conjugated anti-rabbit (Invitrogen). HRP-peroxidase chemiluminescent substrate (SuperSignal™ West Pico PLUS, Thermo Fisher Scientific) was used to reveal the antibody signal, using a charge-coupled device (CCD)-camera based imaging system (Cytiva).

### Biophysical analysis of photosynthesis

PSII function was assessed by variable chlorophyll *a* fluorescence kinetics induced via a saturating pulse (7700 μmol photons·m^-2^·s^-1^) on cells that were dark-adapted for 15 min.

Fast chlorophyll *a* fluorescence transients (OJIP) were measured using were evaluated using a MultispeQ device (PhotosynQ Inc) connected to the PhotosynQ platform (http://www.photosynq.org), using the built-in standard sequences. Cells were dark-adapted for 15 min, before 20 μg chlorophyll being filtered onto a glass microfiber paper for the measurements; the fluorescence transients were normalized against the minimal (F_0_) and maximal (F_M_) fluorescence and plotted on a logarithmic time scale.

PSI function was assessed by absorbance kinetics at 705 nm with background correction at 740 nm (P_700_) and ElectroChromic Shift (ECS) of carotenoids was measured at 520 nm with background correction at 546 nm; for P_700_ measurements, the traces were normalized against the intensity signal of cells treated with 20 μM of 3-(3,4-dichlorophenyl)-1,1-dimethylurea (DCMU) and 40 μM of 2,5-dibromo-3-methyl-6-isopropylbenzoquinone (DBMIB). Cells were acclimated for 3 min at each light intensity (110, 210, 480, 1100, 2100, 4300 µmol photons m^−2^ s^−1^), and the measurements used a Joliot-type Spectrophotometer (JTS-Biologic) and a Continuum laser for single turnover saturating flashes, as described on (56); each biological replicate measurement is the result of 3 technical replicate measurements.

CO_2_ assimilation was measured using a LI-COR 6800-18 aquatic chamber, based on (57). Before each measurement, 0.8 OD_750_ equivalents of cells were harvested via centrifugation (1100 g, 5 min, RT) and re-suspended in 15 ml of air-saturated fresh TAP medium supplemented with 0.5 mg carbonic anhydrase. Cells were then acclimated for 6 min at each light intensity (0, 50, 150, 300, 600, 900, 1200, and 1500 µmol photons m^−2^ s^−1^) followed by a logging event, where CO_2_ concentration was measured and a saturating flash was applied to measure chlorophyll *a* fluorescence parameters.

Maximal O_2_ evolution was measured using an needle-Type Microx TX3 Oxygen Microsensor (Pre-Sens, Precision Sensing GmbH), based on (58, 59) with minor modifications. 3.0 ml of culture were collected, supplemented with 0.1 mM of the artificial acceptor 6-dichloro-1,4-benzoquinone (DCBQ) and 0.5 mM of potassium ferricyanide (FeCy), and continuously stirred with a magnetic stir bar in darkness. After 60 s of dark incubation, when a stable baseline was reached, the sample was illuminated with continuous actinic light (about 5000 µmol photons m^−2^ s^−1^); the dissolved oxygen concentration was measured continuously by the manufacturer’s software (TX3v602) for 6 min, at a sampling interval of 1 s. The oxygen sensor was calibrated with two points, that is, air‐saturated water (100 % air saturation) and anoxic water (1 % Na_2_SO_3_) before the measurement. The oxygen evolution rate was determined from the slope of the linear fitting of the original traces during the illumination phase and expressed in μmol O_2_ h^-1^ mg Chl^-1^ units.

### Recombinant protein production

The protocol for production of recombinant proteins was based on (8). The synthetic DNA sequences of *C. reinhardtii lmtp1* (corresponding to the residues R77 to K246 of Cre15.g636050) and *tlp26* (corresponding to the residues A82 to V226 of Cre03.g154600) genes, flanked by complementary *BsaI* restriction sites sequences, were ordered from Twist Bioscience and sub-cloned into the plasmid pNIC28-Bsa4 (Addgene#26103) using GoldenGate cloning technique. The pNIC28-Bsa4 vector contains a 6×His-tag and a TEV protease-cleavage site followed by the suicide gene *sacB* (flanked by *BsaI* restriction sites), which is replaced by our synthetic gene sequences. 20 fmol of vector was used with 20 fmol synthetic DNA for three successive rounds of digestion (*BsaI*, 37°C, 10 min) and ligation (T4 DNA ligase, 16°C, 10 min) in the same mix. TOP10 *E. coli* cells (NEB) were then transformed with the ligation products for vector amplification, screening and sequencing. 50 ng of vector were then used to transform chemically competent *E. coli* BL21(DE3) cells (Invitrogen) for protein expression.

Bacterial cells were cultured at 37 °C in Luria Broth (Lennox) supplemented with 50 µg ml^-1^ of kanamycin. Expression was induced once cells reached an OD_600_ of 0.6-0.8 by addition of 500 μM IPTG, followed by incubation overnight at 17 °C. Cells were collected by centrifugation and re-suspended in lysis buffer (50 mM Sodium-Phosphate buffer, 300 mM NaCl, 10 mM Imidazole, pH 8.0) supplemented with protease inhibitor cocktail (Sigma-Aldrich) and DNase. Lysis buffer was used in the ratio of 1.5 ml buffer per 1 g of wet cell pellet. Re-suspended pellets were stored at - 80 °C.

Cells were disrupted twice by French Press at 1000 psi and clarified by centrifugation at 40000 x *g* for 45 min. Purifications of the His-tagged proteins were performed in two steps, using Ni-charged HisTrap HP (Cytiva) and HiLoad 16/60 Superdex 200 pg columns (Cytiva) on an ÄKTA Pure FPLC protein purification system (Cytiva); the columns were pre-equilibrated with IMAC buffer (50 mM Sodium-Phosphate, 300 mM NaCl, 10 mM Imidazole, pH 8.0) and gel filtration buffer (20 mM MES, 150 mM NaCl, pH 6.5), respectively. The 0.45 µM filtered lysates were loaded onto the HisTrap HP column and washed with IMAC buffer. Bound proteins were eluted with IMAC buffer containing 250 mM imidazole and cleavage of the 6xHis-tag were done overnight at 4 ºC, using recombinant TEV protease at a 1:20 (TEV:target protein) ratio by dialysis against IMAC buffer without imidazole. Cleaved proteins were separated from the tag and the TEV protease by a second run on Histrap HP column and loaded onto the pre-equilibrated gel filtration column. Fractions containing the target proteins were pooled and concentrated using a VIVASPIN centrifugal filter device with a cut-off size of 10 kDa. Protein purity was confirmed by SDS– polyacrylamide gel electrophoresis and protein samples were flash frozen in liquid nitrogen and stored at -80 °C.

### Crystallization, data collection, structure determination and validation

Native crystals were obtained by the “sitting drop vapour” diffusion method in a 96-well plate for both CrLMTP1 and CrTLP26. 0.5 µl of protein solution (at a concentration of 6.3 mg ml^-1^ and 10.2 mg ml^-1^, respectively) in gel filtration buffer (20 mM MES, 150 mM NaCl, pH 6.5) was mixed with 0.5 µl of “well solution”, consisting of 0.2 M Sodium phosphate dibasic dihydrate at pH 9.1 and 20 % (w/v) Polyethylene glycol 3,350 for CrLMTP1, and 0.2 M Ammonium sulfate, 0.1 M BIS-TRIS pH 5.5, and 25 % (w/v) Polyethylene glycol 3,350 for CrTLP26. In both cases, the plate was incubated at 20 °C and crystals were obtained after 12-14 days. Crystals were quickly transferred to cryogenic solution containing “well solution” supplemented with 20 % glycerol, and flash frozen in liquid nitrogen.

For both proteins, native data were collected to 1.9 Å on beamline ID30B at the ESRF Synchrotron facility (France). Data integration and scaling were performed with Grenades fastproc or AutoPROC pipeline on the beamline. The structures were solved by molecular replacement with PHASER and MrBUMP, respectively in the CCP4 suite (60).

Models were generated by successive rounds of building and refinement with Coot and Refmac5 and/or Phenix (61–63). Structure validations were performed using the Molprobity server (http://molprobity.biochem.duke.edu/) and Procheck from the CCP4 suite (60). Coordinates and structure factors have been deposited in the Protein Data Bank under PDB ID 9H70 (CrLMTP1) and PDB ID 9TNJ (CrTLP26); information regarding crystal data, data-collection, and refinement statistics can be found in Table S2.

### *Thermal unfolding experiments via* nano-differential scanning fluorimetry *(nanoDSF)*

For thermal unfolding experiments, proteins were diluted to a final concentration of 10 µM and incubated with 5 molar excess of each divalent metal. In redox experiments, proteins were first incubated for 4 h at 25 ºC with freshly prepared 2 mM H_2_O_2_ or 2 mM tris(2-carboxyethyl)phosphine (TCEP). For each condition, 10 µl of sample per capillary were required for one thermal unfolding profile. The samples were loaded into Prometheus capillaries (NanoTemper Technologies) and experiments were carried out using the Prometheus NT.48. The temperature ramp was set to an increase of 1 °C/min in a range from 20 °C to 90 °C. Protein unfolding was measured by detecting the temperature-dependent change in tryptophan fluorescence at emission wavelengths of 330 and 350 nm. For calculation of Melting Temperature (Tm), the first derivative data of the F_350_/F_330_ fluorescence ratio unfolding curves were used.

### Phylogenetic analysis

A phylogenetic tree of the thylakoid acid phosphatase family was built from sequences of the PANTHER entry PTHR35514 / Thylakoid_lumenal_15kDa-like, some of which are also classified in the InterPro entry IPR007621 / TPM_dom family. A set of homologous amino-acid sequences representing the major taxa were selected in the dbUniProtKB reference proteomes and Swiss-Prot databases (July 2024), including eukaryotic lineages from the *Streptophyta, Chlorophyta, Rhodophyta*, and the *Stramenopila*-*Alveolata*-*Rhizaria* group. A multiple sequence alignment was performed with MAFFT v7.49 (64) using the L-INS-i strategy, and filtered for saturation with BMGE v1.12 (65) using the BLOSUM30 matrix and removing sites with > 50% gaps, to get a final alignment containing 189 sites and 51 sequences. The tree was built using the Maximum-Likelihood (ML) algorithm with IQ-TREE (66) and the Q.pfam+I+R4 substitution model selected by ModelFinder (67) with the Bayesian Information Criterion (BIC). The statistical support of the branches was estimated using a (UFBoot) approximation (68) implemented in IQ-TREE with 1000 replicates.

### Statistical analysis

When applicable, data is presented as Average ± Standard error of the mean. Statistical significance was determined using one-way ANOVA followed by Dunnett multiple comparison posttests (Prism 10; GraphPad Software). Changes were considered statistically significant at p < 0.05 and presented as: p < 0.05, 1 symbol (*); p < 0.01, 2 symbols (**); p < 0.001, 3 symbols (***).

## Acknowledgments

We thank Géraldine Brandelet and Clara Amselem of the ProteinTec platform of the BIAM for their help with protein production and crystallization. We also warmly thank Bernard Genty, Fabrizio Iacono, Mateusz Szwalec, Anja Krieger-Liszkay, Stefano Caffarri, Franck Chauvat, Pathomchai Dindaeng, and Chloé Maurin for helpful discussions and their involvement in this project.

We are grateful to the INRAE MIGALE bioinformatics platform (https://migale.inrae.fr/) for providing computational resources.

This work was supported by Agence Nationale de la Recherche (ANR) funding awarded to XJ (“ChloroPaths”, ANR-14-CE05-0041-01; “RevelOrg”, ANR-20-CE20-0006) and overheads from PEPR FairCarboN France 2030.

## Accession numbers

Sequence data for the proteins referred in this article can be found in Uniprot under following accession numbers: LMTP1: A8JDW2_CHLRE; TLP26: A0A2K3DVX2_CHLRE; Sll1071: P73281_SYNY3; AT5G52970: TL15B_ARATH; CrTLP18.3: A8JH60_CHLRE. The coordinates of the recombinant CrLMTP1 and CrTLP26 crystal structures were deposited on the RCSB Protein Data Bank (RCSB PDB) under the PDB ID 9H70 and 9TNJ, respectively.

## List of figures and tables

*Figure S1:* Maximum likelihood tree of the TPM-domain family (PANTHER entry PTHR35514 / Thylakoid_lumenal_15kDa-like) showing its distribution in green and red algae, land plants, and *Cyanobacteriota*.

*Figure S2:* Alignment of the mature protein sequences of CrLMTP1, AtLMTP1, Sll1071, CrTLP26, and CrTLP18.3.

*Figure S3:* CrLMTP1 and CrTLP26 are chloroplast-targeted proteins with different predicted secondary structures.

*Figure S4:* Generation of *C. reinhardtii* single *lmpt1* and *tlp26* mutants via CRISPR/Cas9 and their phenotypical analysis during growth in mixotrophic conditions under low (20 µmol photons m^-2^ s^-1^) or moderate high (250 µmol photons m^-2^ s^-1^) light intensities.

*Figure S5:* Generation of *C. reinhardtii lmpt1 tlp26* mutants via genetic crossing of the single *lmtp1* (mt-) and *tlp26* (mt+) mutants.

*Figure S6:* Immunoblots used to prepare the densitometry analysis presented on Figure 3c.

*Figure S7:* Immunoblot analysis of cells grown under photoinhibitory light intensities (650 μmol photons m^-2^ s^-1^) in the absence or presence of chloroplast translation/elongation inhibitors (Lincomycin, chloramphenicol).

*Figure S8:* Biological replicates of the measurements presented in Figure 4.

*Figure S9:* Metal binding prediction to CrLMTP1.

*Table S1:* Conserved genes that have evolved with oxygenic photosynthesis.

*Table S2:* Crystal data, data-collection and refinement statistics.

