## Supplemental Information Figures and tables for "Structural and functional characterization of two conserved lumenal TPM-domain proteins involved in the maturation of Photosystem II"

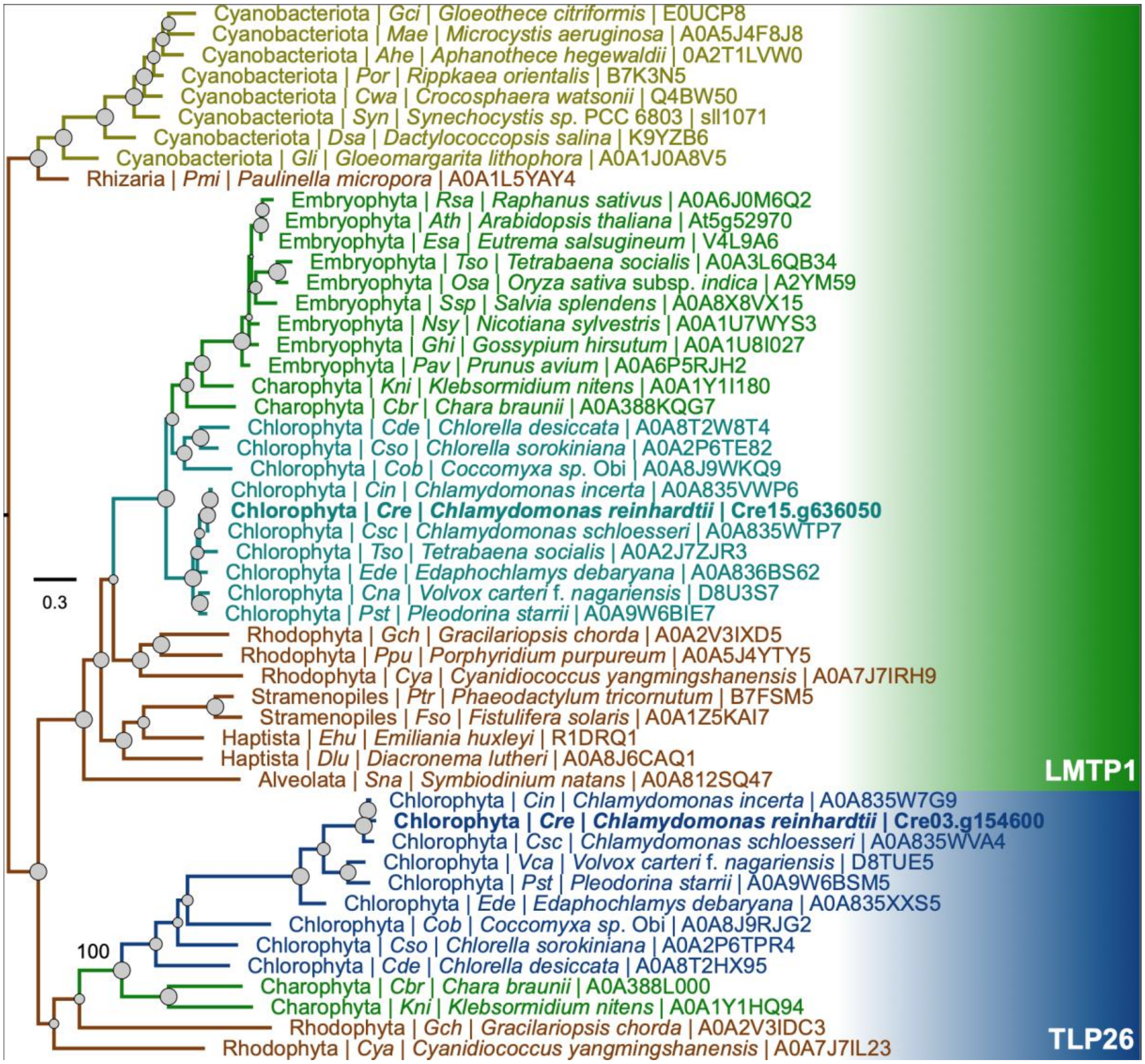


**Fig. S1. Maximum likelihood tree of the TPM-domain family (PANTHER entry PTHR35514 / Thylakoid_lumenal_15kDa-like) showing its distribution in green and red algae, land plants, and *Cyanobacteriota*.** Sequence names are associated with their primary accession numbers in the UniProt database. The tree was rooted with *Cyanobacteriota* since the protein in eukaryotes was likely inherited from the endosymbiotic ancestor that gave rise to the chloroplast. CrLMTP1 homologs in aquatic eukaryotic species and several *Streptophyta* species (including *Charophyta* and *Embryophyta*) do form a monophyletic group, distinct from CrTLP26 homologs found in the same aquatic eukaryotic species, but not in land plants. Given the functional differences, the molecular weight, the poor sequence similarities and the tree topology, we propose to rename these clusters as **L**umenal **M**n-binding **T**PM-domain **P**rotein **1** (LMTP1), and **T**hylakoid **L**umenal **P**rotein **26** kDa (TLP26), respectively. The tree is drawn to scale, and the scale bar represents the number of substitutions per site. Tree topology was tested using an ultrafast bootstrap approximation approach with 1000 replicates; gray circles represent bootstrap values and are drawn to scale.

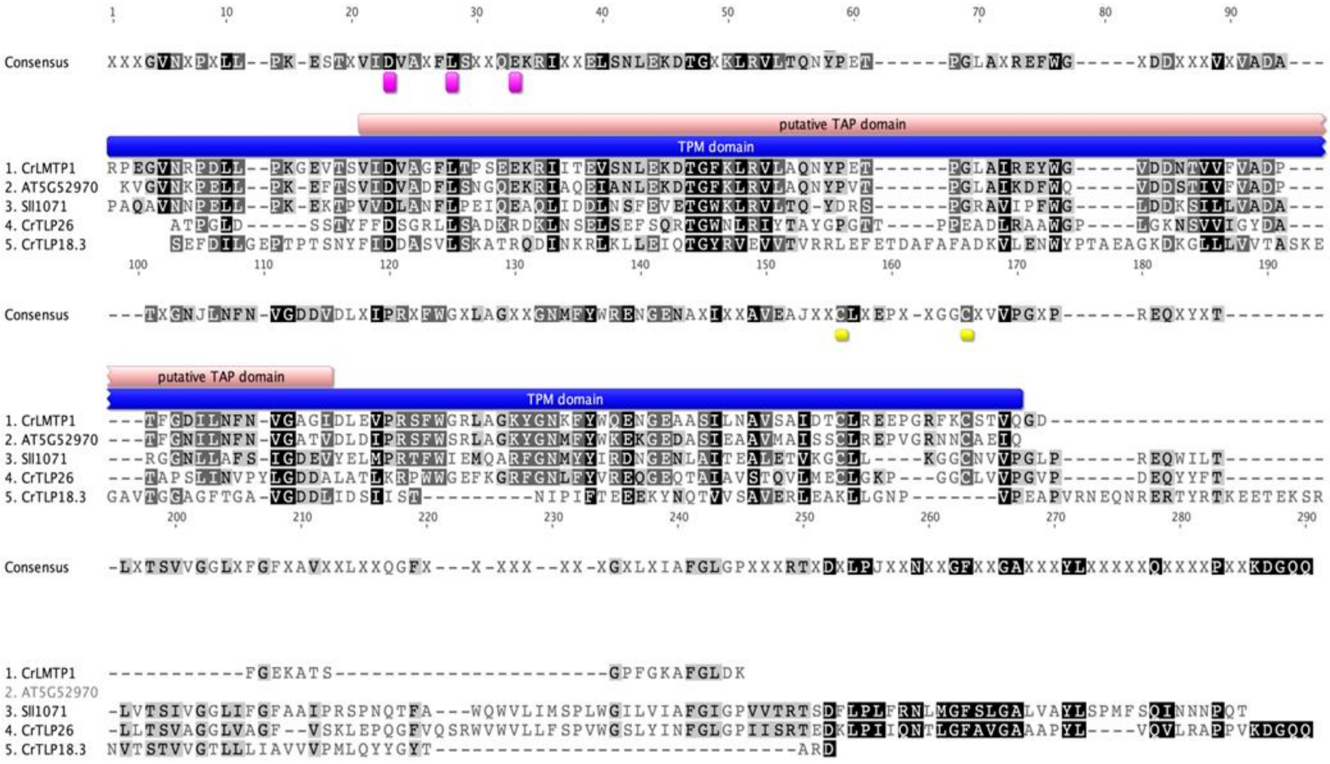

**Fig. S2. Alignment of the mature protein sequences of CrLMTP1, AtLMTP1, Sll1071, CrTLP26, and CrTLP18.3.** Mature protein sequences (i.e., without predicted transit peptides) of LMTP1 from *C. reinhardtii* and *A. thaliana*, Sll1071 from *Synechocystis*, and TLP26 and TLP18.3 from *C. reinhardtii* were aligned using MAFFT with default parameters and the BLOSUM45 matrix. The pink box marks the putative "Thylakoid Acidic Phosphatase" (TAP) domain of the proteins within the TPM fold (marked with a blue rectangle), the purple rectangles mark the amino acids involved in the metal coordination of recombinant CrLMTP1 (see Figure 1c), and the yellow rectangles mark the cysteines forming the intramolecular disulfide bond in both recombinant CrLMTP1 and CrTLP26 (see Figures 1a and 1b).

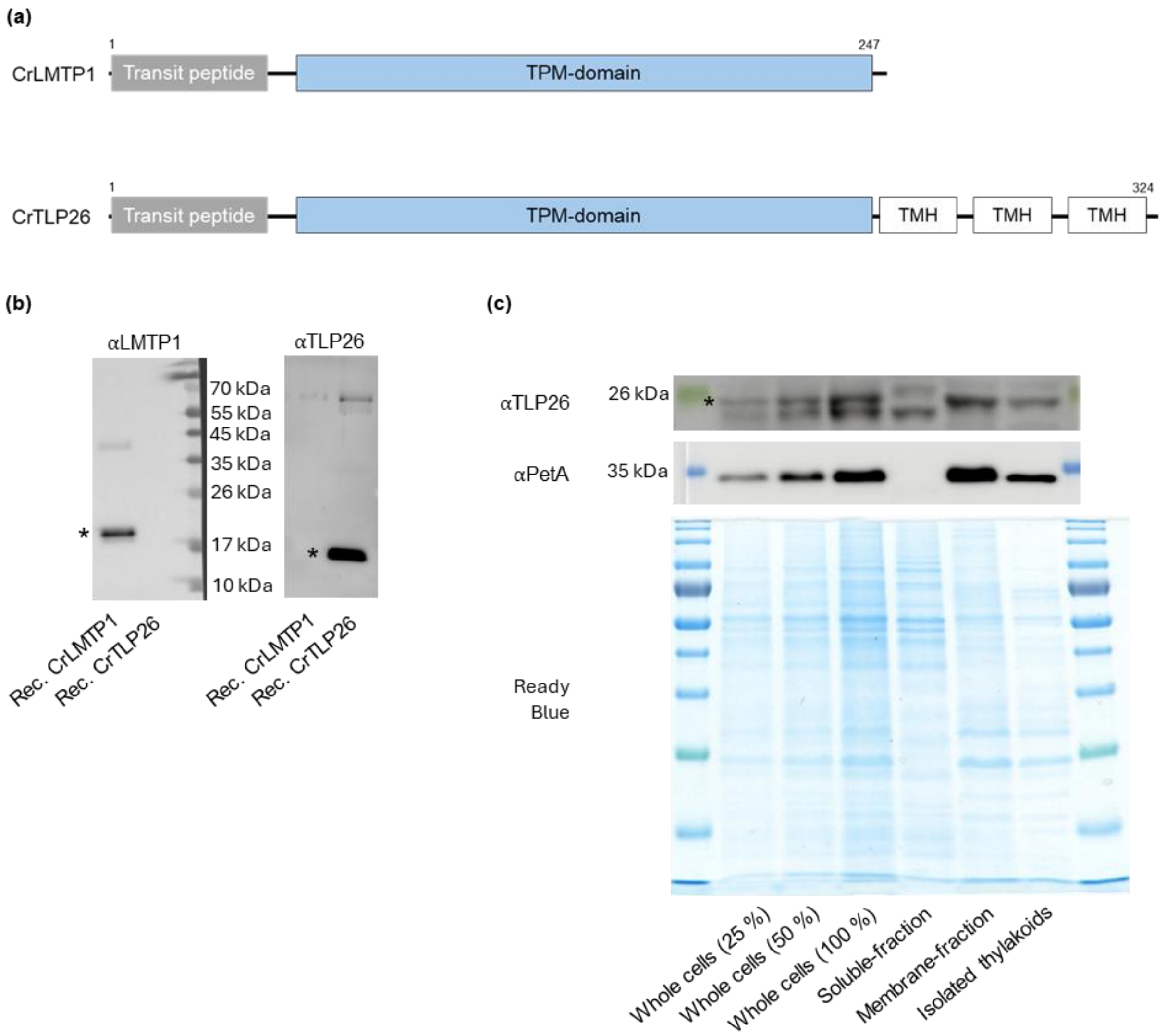


**Fig. S3. CrLMTP1 and CrTLP26 are chloroplast-targeted proteins with different predicted secondary structures. (a)** Predicted secondary structure of both LMTP1 and TLP26 paralogues from *C. reinhardtii*: the N-terminal chloroplast transit peptide is followed by the soluble TPM-domain fold; at the C-terminus of the protein, CrTLP26 contains 3 transmembrane helices that are not present in CrLMTP1. **(b)** Immunoblot analysis of recombinant CrLMTP1 and truncated (i.e. only soluble TPM domain) CrTLP26 proteins, showing that the primary antibodies recognize their specific recombinant protein. **(c)** Immunoblot analysis of whole cell, soluble, membrane-bound, and thylakoid protein extracts from WT *C. reinhardtii* cells, confirming the predicted structure and localization of CrTLP26; PetA was used as a marker for membrane-bound, chloroplastic proteins; 100 % corresponds to 2 µg protein; * indicates the expected protein migration band.

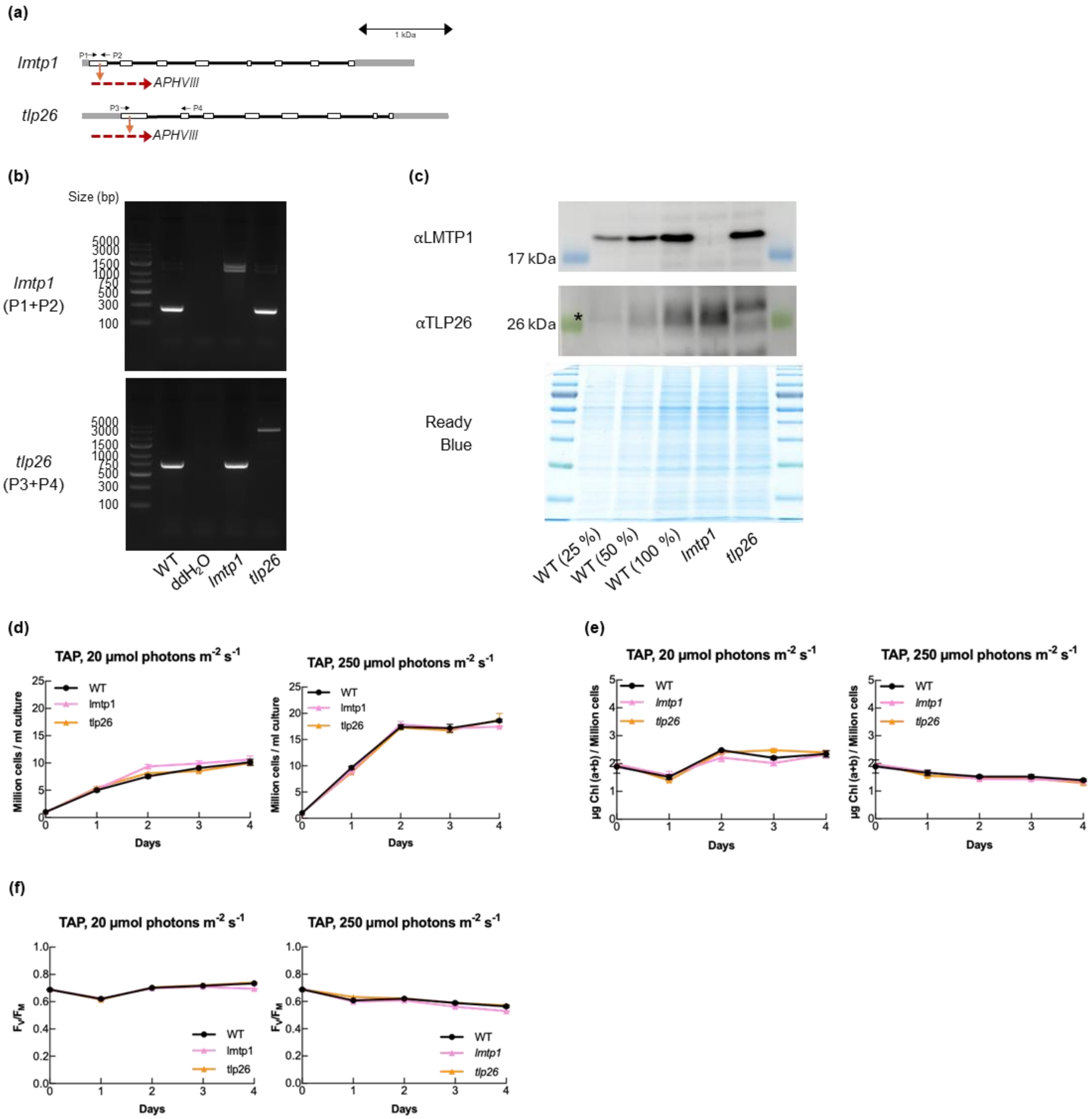


**Fig. S4. Generation of *C. reinhardtii* single *lmpt1* and *tlp26* mutants via CRISPR/Cas9 and their phenotypical analysis during growth in mixotrophic conditions under low (20 µmol photons m$^{-2}$ s$^{-1}$) or moderate high (250 µmol photons m$^{-2}$ s$^{-1}$) light intensities. (a)** Genetic map of Chlamydomonas *lmtp1* (*Cre15.g636050*) and *tlp26* (*Cre03.g154600*), with the prediction mutation site marked by an orange arrow, and with the inserted fragment represented by a red arrow; promoters and terminators are represented as grey boxes, exons as white boxes, and introns as a black line. **(b)** Confirmation of the mutations in the generated strains via PCR, using the primers shown in **(a)** (for the primer sequences and predicted amplification sizes, see the Materials and Methods section of the manuscript). **(c)** Immunoblot analysis of different *C. reinhardtii* strains, showing that the CRISPR/Cas9 mutants cannot accumulate LMTP1 or TLP26; 100 % corresponds to 2 µg protein; * indicates the expected protein migration band. **(d)** Cell density. **(e)** Chlorophyll (*a*+*b*) content per million cells. **(f)** Maximum PSII quantum yield (Fv/Fm). Data in panels **(d)**, **(e)**, and **(f)** is represented as mean ± standard error of the mean (n = 3 replicates).

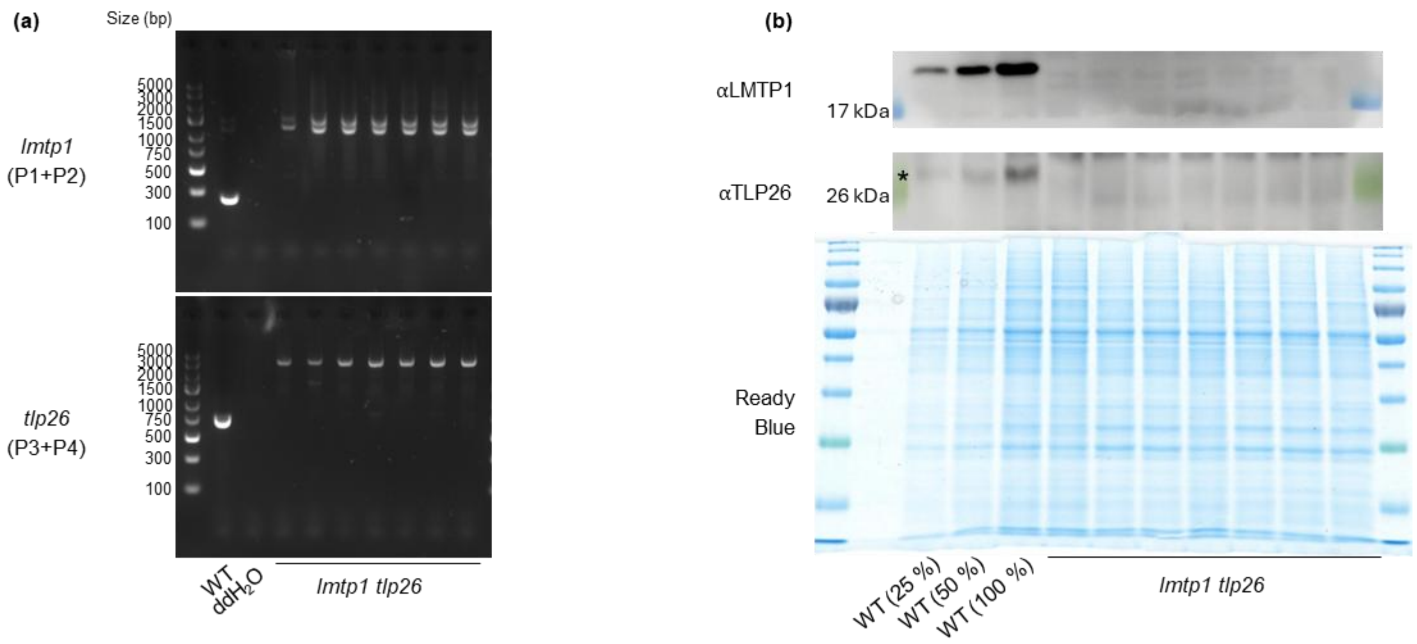


**Fig. S5. Generation of *C. reinhardtii lmpt1 tlp26* mutants via genetic crossing of the single *lmtp1* (mt-) and *tlp26* (mt+) mutants. (a)** Confirmation of the presence of the double mutation via PCR screening. **(b)** Immunoblot analysis of different *C. reinhardtii* strains, showing that the double mutants do not accumulate neither CrLMTP1 nor CrTLP26; 100 % corresponds to 2 µg protein; * indicates the expected protein migration band.

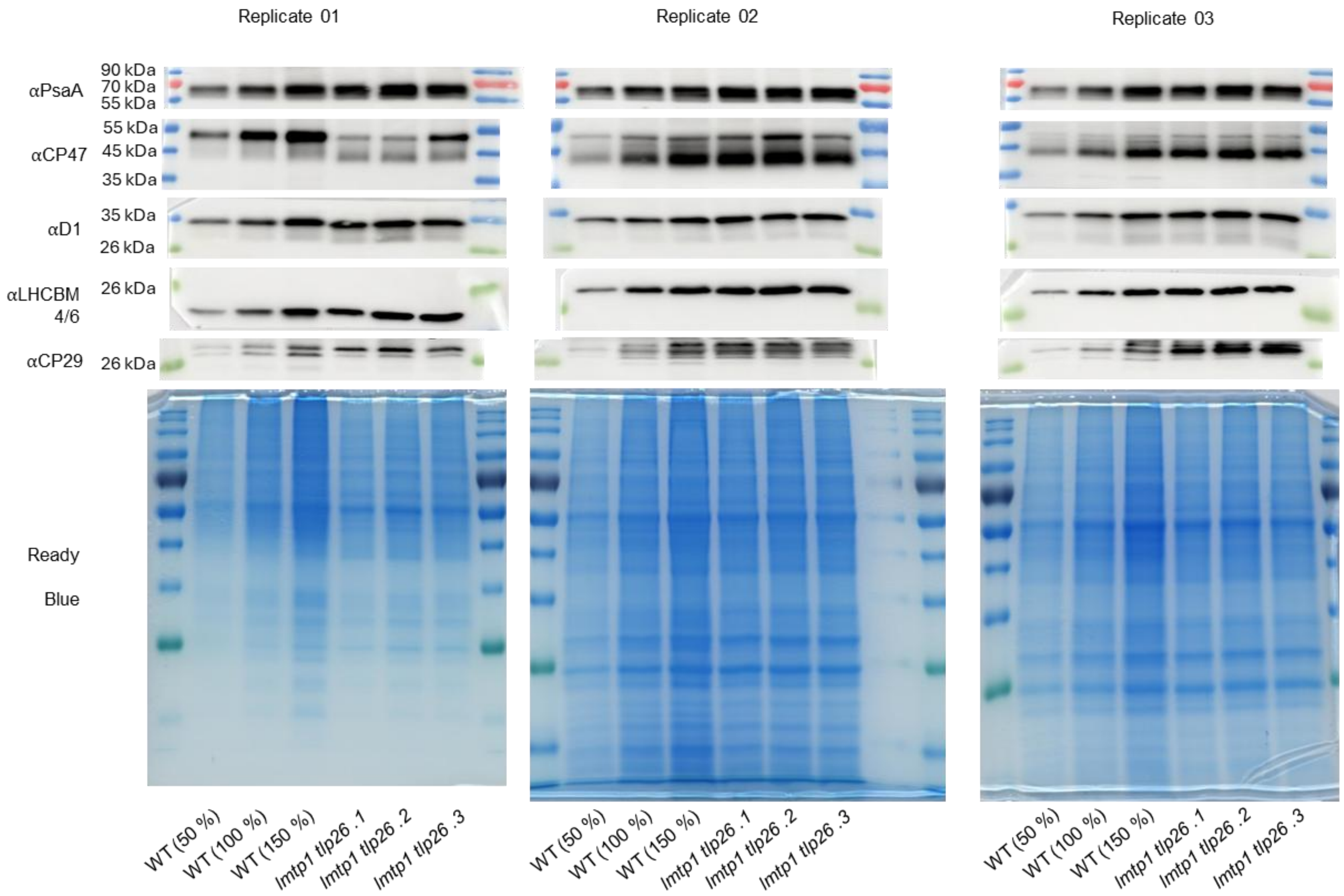


**Fig. S6. Immunoblots used to prepare the densitometry analysis presented on Figure 3c.** Blots were loaded based on total protein quantification, confirmed via Ready Blue staining; 100 % corresponds to 2 μg protein.

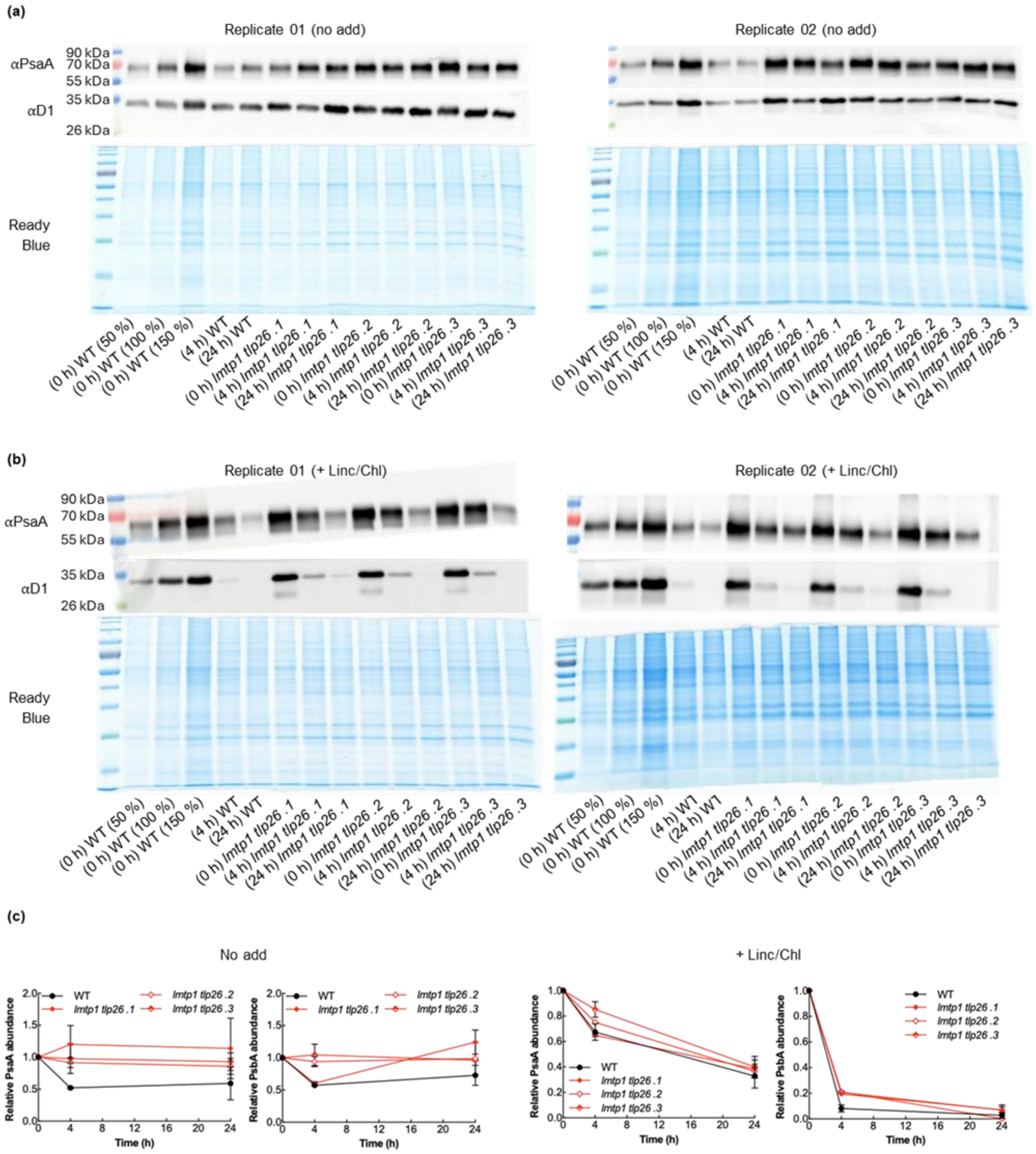


**Fig. S7. Immunoblot analysis of cells grown under photoinhibitory light intensities (650 µmol photons $m^{-2}$ $s^{-1}$) in the absence or presence of chloroplast translation/elongation inhibitors (Lincomycin, chloramphenicol). (a)** Immunoblots against the chloroplast-encoded proteins that are significantly more abundant in *lmtp1 tlp26* lines (see Figures 3c and S6), in the absence of inhibitors. **(b)** Immunoblots against the chloroplast-encoded proteins in the presence of inhibitors. Blots were loaded based on total protein quantification, confirmed via Ready Blue staining; 100 % corresponds to 2 µg protein. **(c)** Densitometry analysis of the immunoblots shown in **(a)** and in **(b)**; data is normalized for the intensity signal of the first time-point of each strain and presented as mean ± standard error of the mean.

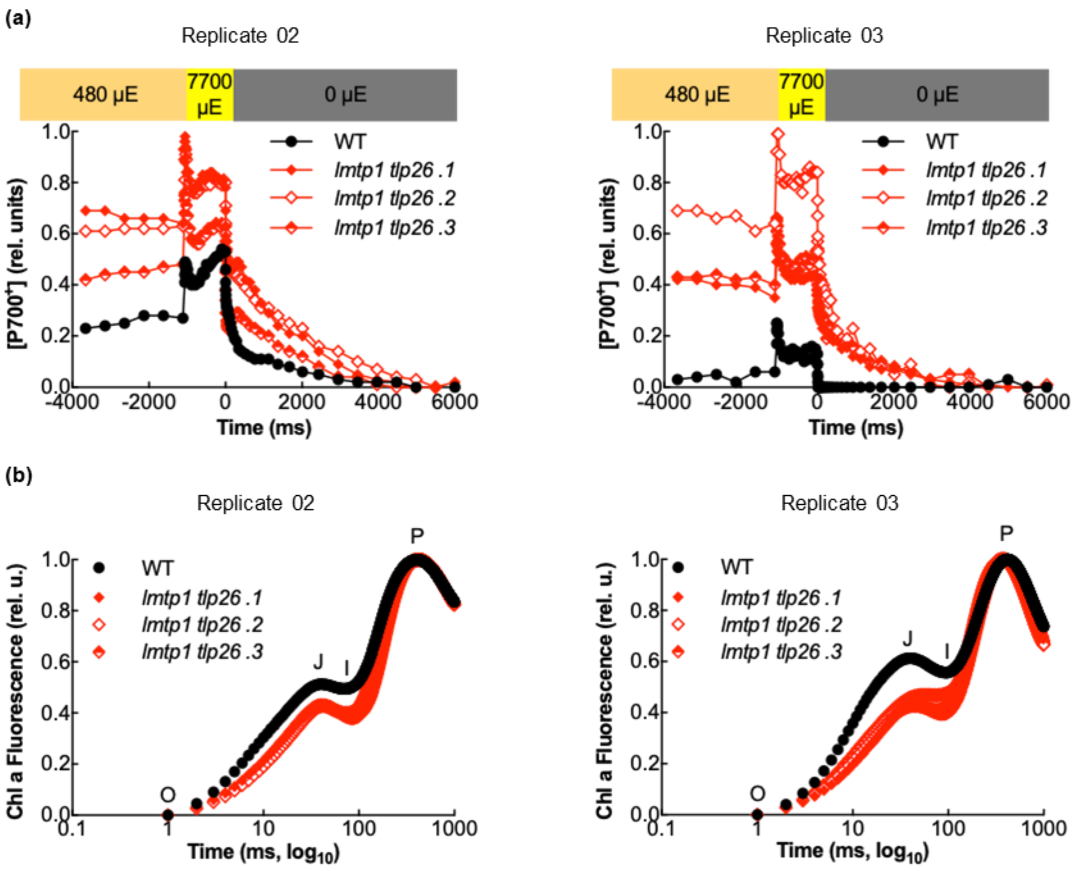


**Fig. S8. Biological replicates of the measurements presented in Figure 4. (a)** $P_{700}$ redox measurements. **(b)** OJIP traces.

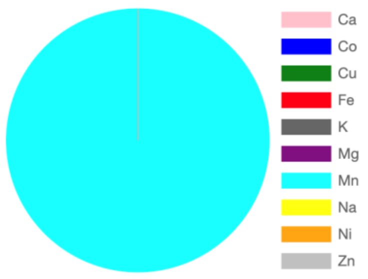


Metal binding predictions at a cutoff of 0.5, to view the scores download the results.

Show 10 entries Search:

| ID | MetalBinding | Ca | Co | Cu | Fe | K | Mg | Mn | Na | Ni | Zn |
|---|---|---|---|---|---|---|---|---|---|---|---|
| LMTP1 | No | No | No | No | No | No | No | Yes | No | No | No |

Showing 1 to 1 of 1 entries Previous 1 Next

**Fig. S9. Metal binding prediction to CrLMTP1.** The full protein sequence of CrLMTP1 (Cre15.g636050) was used as template to test metal interaction probabilities, using the *MeBiPred* server, showing that from all tested divalent metals $Mn^{2+}$ is the likely cofactor of this protein; screenshot taken directly from the server results webpage.

**Tables**

**Table S1. Conserved genes that have evolved with oxygenic photosynthesis.** This table contains a subset of genes that are present in all oxygenic photosynthetic organisms and absent from cyanobacterial accessions that have lost Photosystem II (PSII).

| ***Gene (Synechocystis)*** | ***Annotation*** |
|---|---|
| *slr0906* | psbB (CP47) |
| *sll0851* | psbC (CP43) |
| *ssr3451* | psbE ($\alpha$-$b_{559}$) |
| *smr0006* | psbF ($\beta$-$b_{559}$) |
| *ssl2598* | psbH |
| *sll0427* | psbO |
| *sll1418* | psbP |
| ***slr0575*** | **APE1** |
| ***sll1071*** | **Sll1071 (CPLD31 / TLP15.2)** |

**Table S2. Crystal data, data-collection and refinement statistics.** Values in parentheses correspond to the highest resolution shell.

| DATA COLLECTION (PDB) | | |
|---|---|---|
| **DATA SET NAME** | LMTP1 | TLP26 |
| **PDB ID** | 9H70 | 9TNJ |
| **WAVELENGTH** | 1.77 Å | 1.77 Å |
| **BEAMLINE** | ID30B | 1D30B |
| **TEMPERATURE** | 100 K | 100 K |
| **SPACE GROUP** | P32 | C2 |
| **UNIT-CELL PARAMETERS** | | |
| **A, B, C (Å)** | 51,6, 51,6, 52,5 | 111.8, 43.9, 58.6 |
| **A, B, G (°)** | 90, 90, 120 | 90, 99.3, 90 |
| **RESOLUTION RANGE (Å)** | 25.8 – 1.9 | 97.9 – 1.9 |
| **HIGH RESOLUTION RANGE (Å)** | 2.0 – 1.9 | 1.9 – 1.9 |
| **OBSERVED REFLECTIONS** | 29683 | 22754 |
| **NO. OF UNIQUE REFLECTIONS** | 1475 | 1101 |
| **COMPLETENESS (%)** | 97.2 (74.5) | 99.7 (98.7) |
| **CC1/2** | 100 (0) | 100 (40) |
| **<I/S(I)>** | 21.9 (4.5) | 6.1 (1.0) |
| $R_{MERGE}$**(%)** | 9.5 (19.0) | 14.5 (116.2) |
| **REFINEMENT** | | |
| **RESOLUTION RANGE (Å)** | 44.7 – 1.42 | 57.9 - 1.9 |
| $R_{WORKT}$ / $R_{FREE}$ | 18,12 / 0.21 | 20.11 / 24.9 |
| **NO. OF NON-H ATOMS:** | | |
| **PROTEIN** | 1339 | 2 312 |
| **WATER** | 134 | 79 |
| **B-FACTORS** | | |
| **PROTEIN** | 18.52 | 30.424 |
| **WATERS** | | |
| **R.M.S. DEVIATION FROM IDEAL GEOMETRY** | | |
| **BOND LENGTHS (Å)** | 0.005 | 0.008 |
| **BOND ANGLES (°)** | 0.764 | 1.7 |
